# NUDC is critical for rod photoreceptor function, maintenance, and survival

**DOI:** 10.1101/2023.11.28.568878

**Authors:** Mary Anne Garner, Meredith G. Hubbard, Evan R. Boitet, Seth T. Hubbard, Anushree Gade, Guoxin Ying, Bryan W. Jones, Wolfgang Baehr, Alecia K. Gross

**Affiliations:** Department of Neurobiology, University of Alabama at Birmingham, Birmingham, Alabama, 35294 USA; Department of Ophthalmology and Visual Sciences, University of Utah, Salt Lake City, Utah, 84132 USA

**Keywords:** NUDC, cofilin1 (CFL1), phosphorylated cofilin (pCFL1), photoreceptor, actin, dynein, retinal degeneration

## Abstract

NUDC (<u>nu</u>clear <u>d</u>istribution protein C) is a mitotic protein involved in nuclear migration and cytokinesis across species. Considered a cytoplasmic dynein (henceforth dynein) cofactor, NUDC was shown to associate with the dynein motor complex during neuronal migration. NUDC is also expressed in postmitotic vertebrate rod photoreceptors where its function is unknown. Here, we examined the role of NUDC in postmitotic rod photoreceptors by studying the consequences of a conditional NUDC knockout in mouse rods (r*NudC*^−/−^). Loss of NUDC in rods led to complete photoreceptor cell death at six weeks of age. By 3 weeks of age, r*NudC*^-/-^ function was diminished, and rhodopsin and mitochondria were mislocalized, consistent with dynein inhibition. Levels of outer segment proteins were reduced, but LIS1 (lissencephaly protein 1), a well-characterized dynein cofactor, was unaffected. Transmission electron microscopy revealed ultrastructural defects within the rods of r*NudC*^−/−^ by 3 weeks of age. We investigated whether NUDC interacts with the actin modulator cofilin 1 (CFL1) and found that in rods, CFL1 is localized in close proximity to NUDC. In addition to its potential role in dynein trafficking within rods, loss of NUDC also resulted in increased levels of phosphorylated CFL1 (pCFL1), which would purportedly prevent depolymerization of actin. Absence of NUDC also induced an inflammatory response in Müller glia and microglia across the neural retina by 3 weeks of age. Taken together, our data illustrate the critical role of NUDC in actin cytoskeletal maintenance and dynein-mediated protein trafficking in a postmitotic rod photoreceptor.

**Significance Statement:** Nuclear distribution protein C (NUDC) has been studied extensively as an essential protein for mitotic cell division. In this study, we discovered its expression and role in the postmitotic rod photoreceptor cell. In the absence of NUDC in mouse rods, we detected functional loss, protein mislocalization, and rapid retinal degeneration consistent with dynein inactivation. In the early phase of retinal degeneration, we observed ultrastructural defects and an upregulation of inflammatory markers suggesting additional, dynein-independent functions of NUDC.

## Introduction

The *nud* (<u>nu</u>clear <u>d</u>istribution) genes, including *nudA* (dynein heavy chain), *nudC*, *nudF* (LIS1), *nudG* (dynein light chain or LC8 and *nudK* encode elements of the dynein/dynactin complex in *Aspergillus nidulans* [1]. NudC was identified as a protein that interacts with microtubules and nuclei [2] and regulates dynein-mediated nuclear migration [3–6]. Loss of NudC is lethal in *Aspergillus* [7] and stalls cell division at the one-cell stage in cultured mammalian cells [8].

NUDC has been investigated in human dividing cells [9–12] and is conserved across phyla [8, 9, 13–17]. NUDC is expressed in the nervous system [17] as well as in migrating neurons in cell culture [18]. Our group found that NUDC is expressed in postmitotic vertebrate rod photoreceptors and is required for rod maintenance. Upon shRNA knockdown of *NudC* in the *Xenopus laevis* retina, rod outer segment (OS) discs overgrew and coincided with retinal degeneration [19]. NUDC also interacts with rhodopsin and may affect rhodopsin trafficking mediated by dynein through the inner segment to the connecting cilium [19].

Disc overgrowth accompanying NUDC depletion suggests NUDC involvement in actin dynamics. Studies of vertebrate photoreceptors have established that actin, located at the distal portion of the connecting cilium, participates in nascent disc elaboration [20–25]. Filamentous actin (F-actin) inhibitors cause OS disc overgrowth in retinal eyecups [26–28]. Additionally, knockdown of the actin polymerization-inducing protein Arp2/3 in rods leads to F-actin loss at the proximal OS and the formation of membrane whorls rather than discs [29]. NUDC stabilizes the actin-modulating protein cofilin 1 (CFL1) [30] to modulate actin branching, as well as the balance between filamentous and globular actins [31]. CFL1 plays a critical role in cytoskeletal remodeling by modulating the quantity and length of F-actin [32]. When phosphorylated by Lim kinase, phosphorylated CFL1 (pCFL1) no longer binds F-actin, purportedly preventing the production of branched or G-actin [33]. Understanding actin dynamics is essential to determine how rod OS discs form, are maintained, and how constituent proteins are trafficked between the rod OS and inner segment (IS).

The role of NUDC in postmitotic cells is incompletely understood. Here, iCre75-induced homologous recombination was used to generate rod-specific *NudC* knockout mice. We show that NUDC loss in postmitotic mouse rods leads to photopic and scotopic dysfunction, morphologic changes in the rod OS, glial inflammation, changes in proteins involved in actin cytoskeletal maintenance and retinal degeneration. The plethora of observed phenotypes highlight the essential role of NUDC in rod photoreceptor function regulating the dynamics of the actin cytoskeletal network and the function of dynein.

## Results

### Conditional knockout of *NudC* in the mouse retina

Previously, our work showed NUDC localization in the mouse, tree shrew, rhesus macaque, and *X. laevis* retina, where NUDC and a mutant form of NUDC, NUDC^L280P^, were localized to the IS and outer nuclear layer (ONL) of photoreceptors [19]. To uncover the role of NUDC in postmitotic rod photoreceptors, we generated a rod-specific *NudC* conditional knockout mouse by creating a mouse line with loxP sites flanking exons 3–7 of the mouse *NudC* gene (*NudC^fl/fl^*) (Fig. 1A-B, S1). By breeding *NudC^fl/fl^* mice with mice expressing Cre under the rod opsin promoter (iCre75) [34], we selectively ablated exons 3—7 of *NudC* in rods (r*NudC*^−/−^) (Fig. 1C). Hereafter, *NudC* rod-specific conditional knockout mice (*NudC^fl/fl^*iCre75^+^) are referred to as r*NudC*^−/−^, and heterozygous mice (*NudC^fl/+^* iCre75^+^) are referred to as r*NudC*^+/−^.

**Figure 1.**
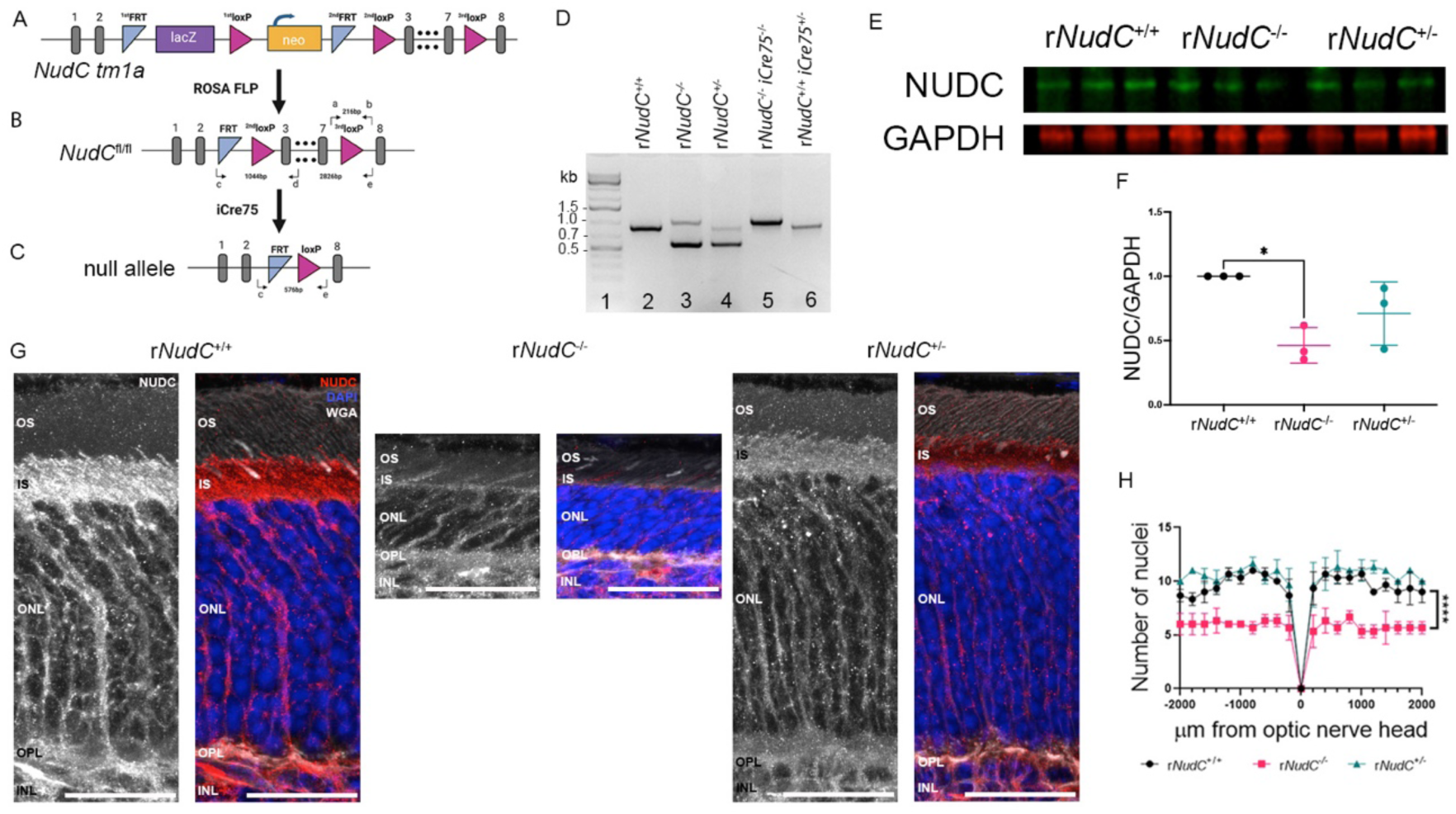
Conditional knockout of *NudC* in the mouse retina. *A*, Schematic diagram of the *Nudc* tm1a clone with gene trap and FRT/loxP sites. *B*, The *Nudc* floxed gene after excision of the gene trap. *C*, The null allele after homologous recombination with *iCre75*. *D*, PCR products for retinal lysates of different genotypes indicated using 3 primers (1: first FRT forward (Fig. 1B, c); 2: Exon 4 reverse (Fig. 1B, d); 3: third FRT LoxP reverse 2, (Fig. 1B, e)). *E*, Western blot of retinal lysates loaded in triplicate indicates a loss of NUDC in r*NudC*^-/-^ but not r*NudC*^+/+^ or r*NudC*^+/-^ at 3 weeks. *F*, Quantification of western blot bands in *E*, n=3 for each genotype. *p<0.05. *G*, Immunohistochemistry of NUDC in greyscale (grey) or color (red) shows staining in the inner segment (IS), outer nuclear layer (ONL), outer plexiform layer (OPL) and inner nuclear layer (INL) of r*NudC*^+/+^. r*NudC*^-/-^ retina exhibits loss of NUDC in the IS and ONL compared with r*NudC*^+/+^ but retains NUDC staining in the INL. Blue, DAPI; Red, NUDC; Grey, wheat germ agglutinin. Scale bar, 20 µm. *H*, Quantification of hematoxylin and eosin (H&E) stain and ONL nuclei counts every 200 µm from the optic nerve head indicates loss of ONL nuclei in r*NudC*^-/-^ compared with r*NudC*^+/+^ and r*NudC*^+/-^ at 3 weeks, n=3 for each genotype. *** p<0.0001. PCR, polymerase chain reaction; RPE, retinal pigmented epithelium; OS, outer segment.

Polymerase chain reaction (PCR) analysis of the five genotypes studied provided predicted amplicons for *NudC* (Fig. 1D). Primers were designed to confirm the presence or absence of floxed exons in retinal DNA. Amplification with primers c and d using *NudC*^+/+^ templates will produce one band of approximately 900 bp (Fig. 1D, lane 2). In the presence of the floxed allele (r*NudC*^-/-^), the presence of FRT and loxP site will produce a higher molecular weight band at 1 kb. In the presence of iCre75, exons 3-7 will be deleted producing an amplicon of 576 bp (Fig. 1D, lanes 3, 4).

In r*NudC*^-/-^ at 3 weeks, there was significantly less total NUDC protein detected in mouse retinal lysates (Fig. 1E and F). There was still NUDC present in the whole retina western blot lysates, and we have previously shown that NUDC localizes not only to the IS and outer nuclear layer of rod photoreceptors but also to the retinal pigmented epithelium (RPE) [19]. Here, using immunofluorescence to stain for NUDC, we show that in r*NudC*^-/-^, while there was diminished NUDC found in the IS and ONL of r*NudC*^-/-^ as compared with r*NudC*^+/+^, NUDC was still present in the inner nuclear layer (INL), inner and outer plexiform layers (IPL and OPL) and retinal ganglion cell (RGC) layer of the same retina (Fig. 1G, middle panels, and Fig. S2). Consistent with the western blotting quantification, there was a noticeable decrease in NUDC stain in the r*NudC*^+/-^ retina compared with r*NudC*^+/+^ (Fig. 1G, right panels).

### Loss of NUDC in rods triggers retinal degeneration as early as 3 weeks postnatal

We next determined whether any change in retinal thickness occurred via histological staining and counting nuclei within the ONL in 200-μm increments from the optic nerve head in 3-week-old animals. Spidergram (number of nuclei measured at various distances from the optic nerve head) analysis of hematoxylin and eosin-stained retinal cryosections (Fig. 1H) revealed a 40% loss of ONL nuclei in r*NudC*^−/−^ mice compared with that in r*NudC^+/+^* and r*NudC*^+/−^ mice as well as that in r*NudC*^−/−^ or r*NudC*^+/−^ control mice (Fig. S3). This difference in retinal thickness across the ONL of the retina can also be seen in the IHC images for NUDC (Fig. 1G).

### NUDC deficiency in rod photoreceptors results in ultrastructural malformations and mislocalized mitochondria

We evaluated the ultrastructure of the remaining rods in 3-week-old mice by transmission electron microscopy (TEM). TEM in 3-week-old r*NudC*^−/−^ mice revealed a thinning retina, and while the remaining OSs appeared relatively normal compared with those of r*NudC*^+/+^ mice (Fig. 2A), r*NudC*^−/−^ rod cells frequently had an extended periciliary ridge (Fig. 2B, asterisks), and in some cases, the discs in the r*NudC*^−/−^ retina grew parallel to the axoneme (Fig. 2B, arrowhead). TEM of the retina of 3-week-old r*NudC*^+/−^ mice demonstrated compact nascent discs with morphology comparable to that of r*NudC*^+/+^ mice (Fig. 2C). Lack of NUDC also resulted in mislocalized mitochondria within the IS. In r*NudC*^+/+^ mice, mitochondria were located against the plasma membrane (Fig. 2D), and in knockout r*NudC*^−/−^ mice, mitochondria appeared to be smaller and dispersed throughout the IS (Fig. 2E). At 6 weeks of age, the r*NudC*^+/−^ retina displayed discs similar to those of r*NudC*^+/+^ mice (Fig. S4C). However, at 6 weeks of age, rod photoreceptors in r*NudC*^−/−^ mice had completely degenerated, with no detectable OS region (Fig. S4B).

**Figure 2.**
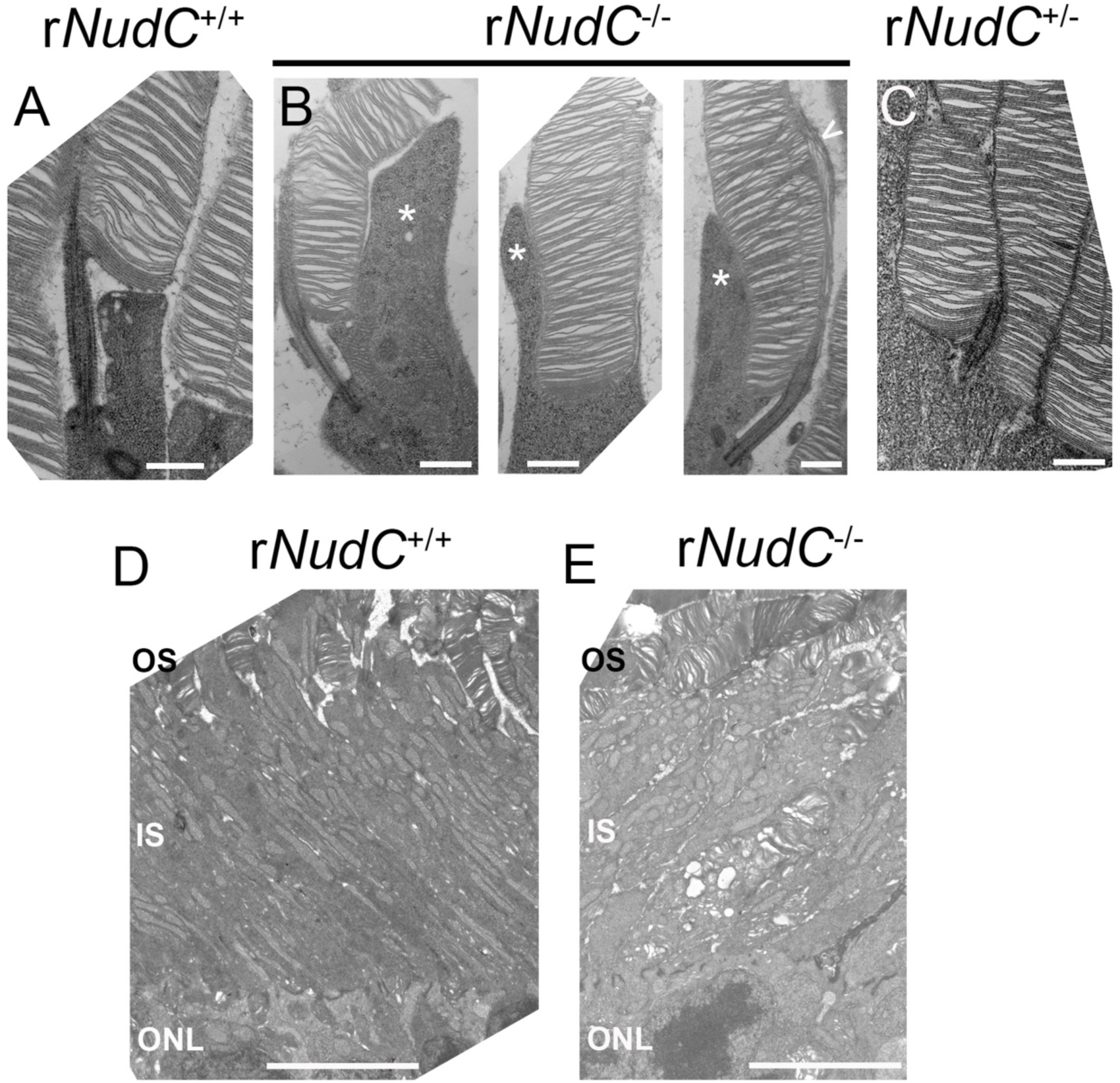
Transmission electron microscopy (TEM). *A-E*, images taken from 3-week-old mouse rod photoreceptors. *A*, *D*, r*NudC*^+/+^; *B*, *E*, r*NudC*^-/-^; *C*, r*NudC*^+/-^. Asterisks in all 3 panels in *B*, regions of extended periciliary ridge; arrowhead, region of abnormal disk overgrowth. *A-C*, scale bar = 600 nm; *D-E*, scale bar = 5 μm. Absence of NUDC rod photoreceptors results in occasional dysmorphic disk membranes, abnormal mitochondria localization, and retinal degeneration.

### Rod function loss in mice lacking *NudC*

We determined whether the loss of NUDC in rods specifically affected retinal function in 3-week-old and 6-week-old mice using electroretinography (ERG). At 3 weeks of age, r*NudC*^+/−^ ERG traces were indistinguishable from those of r*NudC^+/+^* controls, while r*NudC*^−/−^ ERG traces (Fig. 3A) displayed diminished *a*-wave and *b*-wave amplitudes at 0.25 cd-s-m^-2^ and 25 cd-s-m^-2^, with a significant difference in overall light intensities [*p<0.05, **p<0.01] (Fig. 3B). *The a*-wave and *b*-wave implicit times of each genotype were indistinguishable across all light intensities (Fig. 3B).

**Figure 3.**
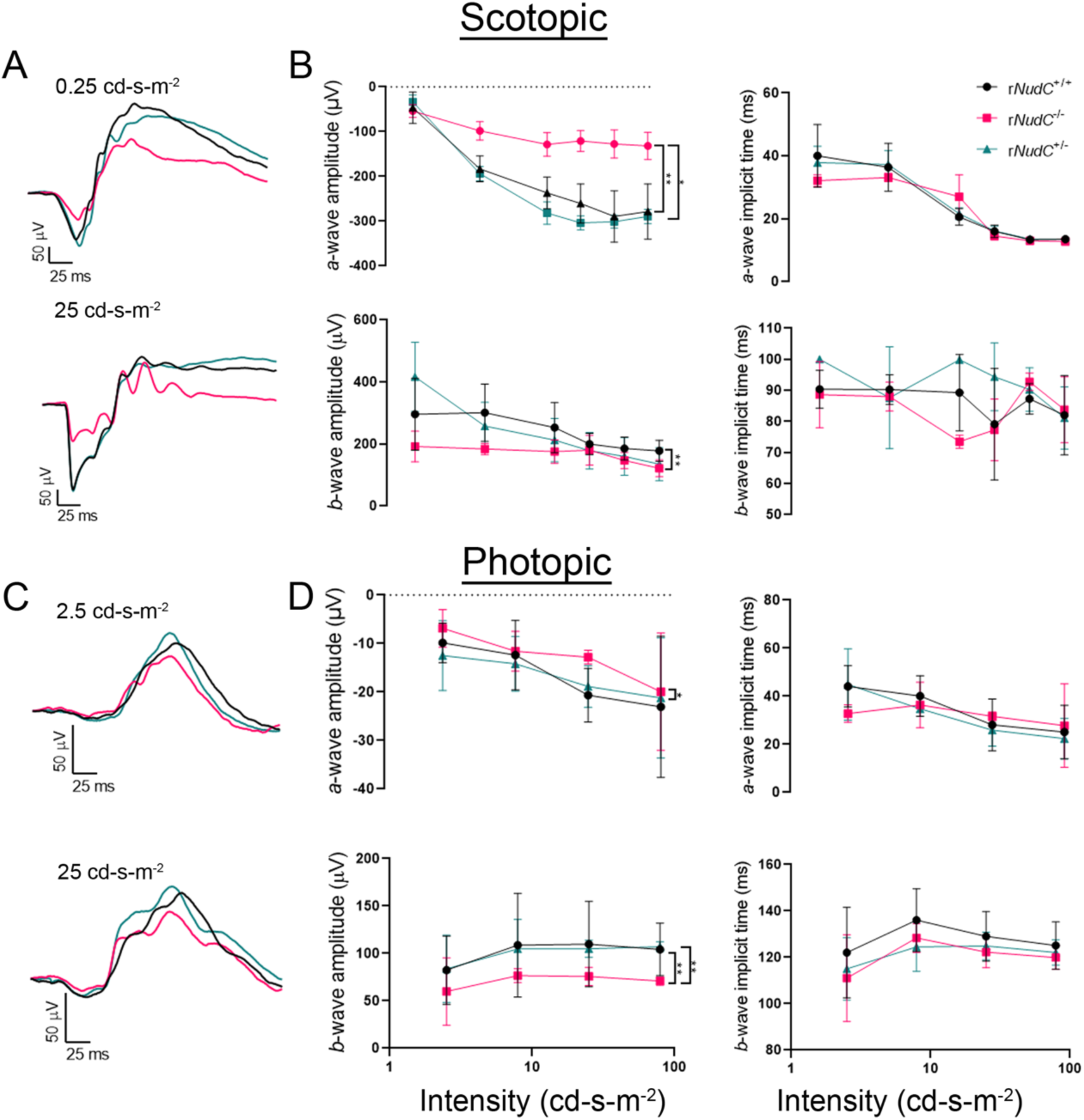
Scotopic and photopic electroretinography. Loss of NUDC affects retinal function in 3-week-old mice. *A*, average electroretinography recordings (n=3) from the lowest (0.25 cd s/m^2^) and highest (25 cd s/m^2^) flash intensities under scotopic conditions for 3-week-old mice. r*NudC*^+/+^, black traces; r*NudC*^-/-^, pink traces; r*NudC*^+/-^, green traces. *B*, summary data of *a*-wave and *b*-wave amplitudes (left panels) and implicit times (right panels) as a function of light intensity from 3-week-old mice. r*NudC*^+/+^, black; r*NudC*^-/-^, pink; r*NudC*^+/-^, green. *p<0.05, **p<0.001. *C*, average electroretinography recordings (n=3) from the lowest (2.5 cd s/m^2^) and highest (25 cd s/m^2^) flash intensities under photopic conditions. r*NudC*^+/+^, black traces; r*NudC*^-/-^, pink traces; r*NudC*^+/-^, green traces. *D*, summary data of *a*-wave and *b*-wave amplitudes and implicit times as a function of light intensity from. r*NudC*^+/+^, black; r*NudC*^-/-^, pink; r*NudC*^+/-^, green. * p<0.05, **p< 0.001.

The *a*-wave and *b*-wave trace amplitudes of 3-week-old r*NudC*^−/−^ and r*NudC^+/+^* mice are displayed under photopic conditions at 2.5 cd-s-m^-2^ and 25 cd-s-m^-2^ (Fig. 3C). There was a significant decrease in both *a*-wave and *b*-wave amplitudes [*p<0.05, **p<0.01, ***p<0.001]; however, no difference in *a*-wave or *b*-wave implicit times was observed (Fig. 3D).

Under scotopic conditions, the floxed *NudC* (r*NudC*^fl/f^ iCre75^−^) and iCre75 positive (r*NudC*^+/+^iCre75^+^) control genotypes showed slight differences, exhibiting a slightly larger *a*-wave amplitude and longer *b*-wave implicit time (Fig. S5A, B). Under photopic conditions, there was no difference in the *a-*wave amplitude, but r*NudC*^+/+^iCre75^+^ mice exhibited a delay in the *a*-wave implicit time. There was a slight difference in the *b*-wave amplitude, and differences in both genotypes were observed for the *b*-wave implicit time compared with that in the r*NudC*^+/+^ control (Fig. S5C, D) [*p<0.05, ***p<0.001].

At 6 weeks of age, ERG traces in the scotopic condition showed no response in r*NudC*^−/−^ mice (Fig. S6A), and the *a*-wave and *b*-wave were diminished at all light intensities [*p<0.05, ** p<0.01] (Fig. S4B). Photopic traces in 6-week-old mice indicated a lack of rod or cone responses in r*NudC*^−/−^ mice (Fig. S6C), a phenotype expected due to the lack of photoreceptors (Fig. 2H). Statistical analysis indicated a significant decrease in the *a*-wave amplitude and implicit time at all flash intensities for r*NudC*^−/−^ mice compared with r*NudC^+/+^*mice [*p<0.05, **p<0.01] (Fig. S6D). A marked difference was also observed in the *b*-wave amplitude and implicit time in r*NudC*^+/−^ mice compared with r*NudC^+/+^* mice [**p<0.01, ***p<0.0005] (Fig. S4D). Functional losses were not observed in r*NudC*^fl/fl^iCre75^−/−^ or r*NudC*^+/+^iCre75^+/−^ control mice (Fig. S7).

### Loss of rod photoreceptors is concurrent with loss of rod OS proteins

Western blotting at 3 weeks of age showed that rod-specific *NudC* knockout resulted in a statistically significant loss of proteins that are normally trafficked to the OS or between the IS and OS of rod photoreceptors, including rhodopsin (RHO) and transducin (GNAT1), compared with *rNudC*^+/+^ mice (Fig. 4A, B). Proteins normally located in the IS (LIS1 and RAB11A) remained unchanged in each genotype (Fig. 4A, B).

**Figure 4.**
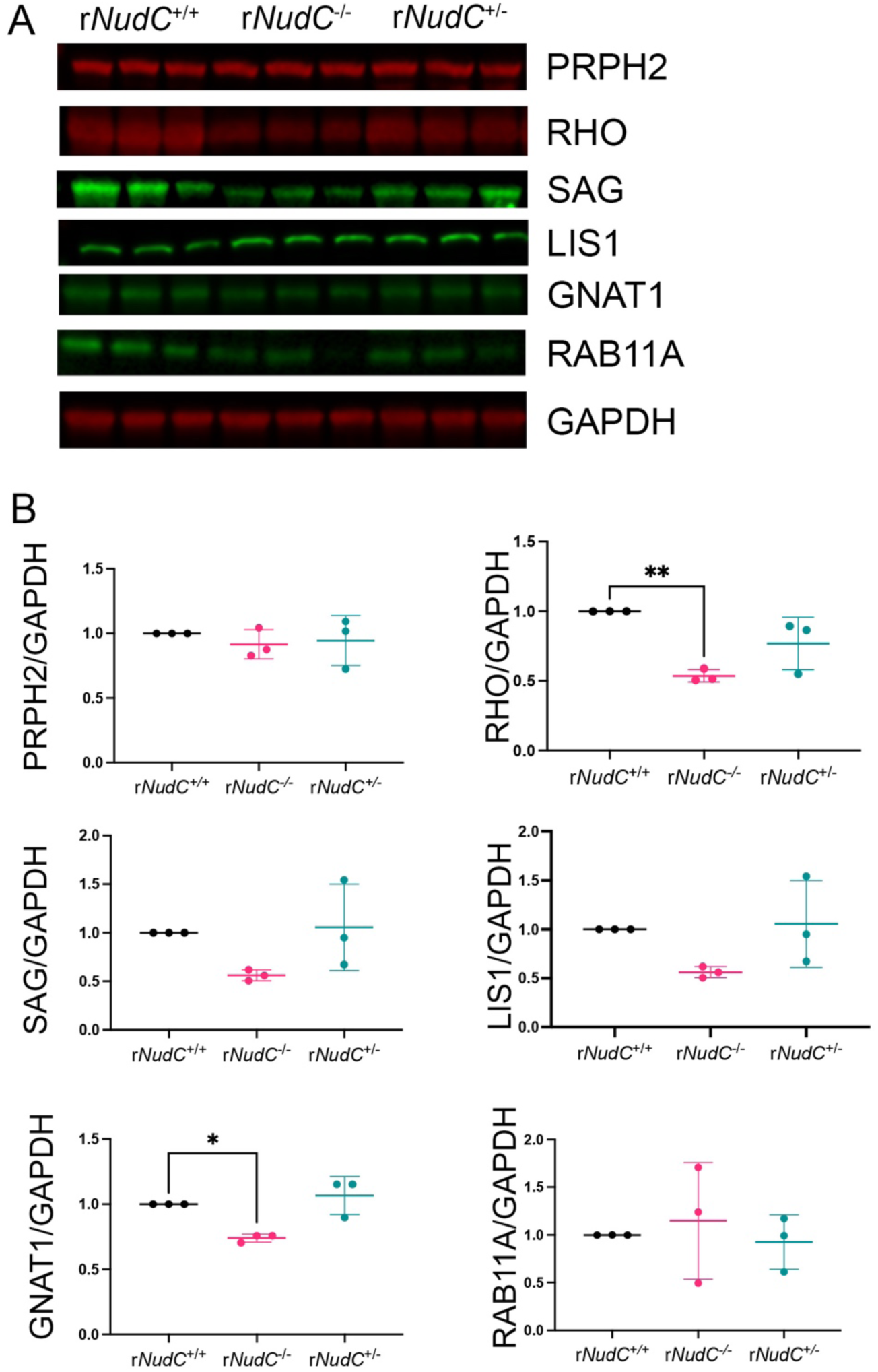
Protein levels affected by the absence of *NudC*. *A*, representative western blots of 3-week-old mouse retina, loaded in triplicate and probed for outer segment (OS) proteins peripherin (PRPH2) and rhodopsin (RHO), inner segment (IS) proteins LIS1 and RAB11A, and proteins that are translocated between the IS and OS, arrestin (SAG), and transducin (GNAT1), depending on light intensity. *B*, Quantification of the western blots described in *A* revealed a significant loss of RHO and GNAT1 in r*NudC*^-/-^ compared with r*NudC*^+/+^. n=3 for each genotype, *p<0.05, **p<0.01.

Immunofluorescence staining revealed mislocalization of RHO in r*NudC*^−/−^ mice compared with the normal distribution of RHO only in the OS in the *rNudC*^+/+^ mouse retina (Fig. 5A). Interestingly, mild mislocalization (Fig. 5A, arrow) of RHO was observed in the r*NudC*^+/−^mouse retina at 3 weeks of age, prior to any retinal degeneration. RHO mislocalization also occurs in 2-week-old r*NudC*^-/-^ mice as well (Fig. S8A). By 3 weeks, the loss of NUDC resulted in several apoptotic TUNEL-positive cells (Fig. 5B, arrows) in the ONL of the r*NudC*^−/−^ retina, while the *rNudC*^+/+^ and r*NudC*^+/−^retinas lacked TUNEL-positive staining (Fig. 5B).

**Figure 5.**
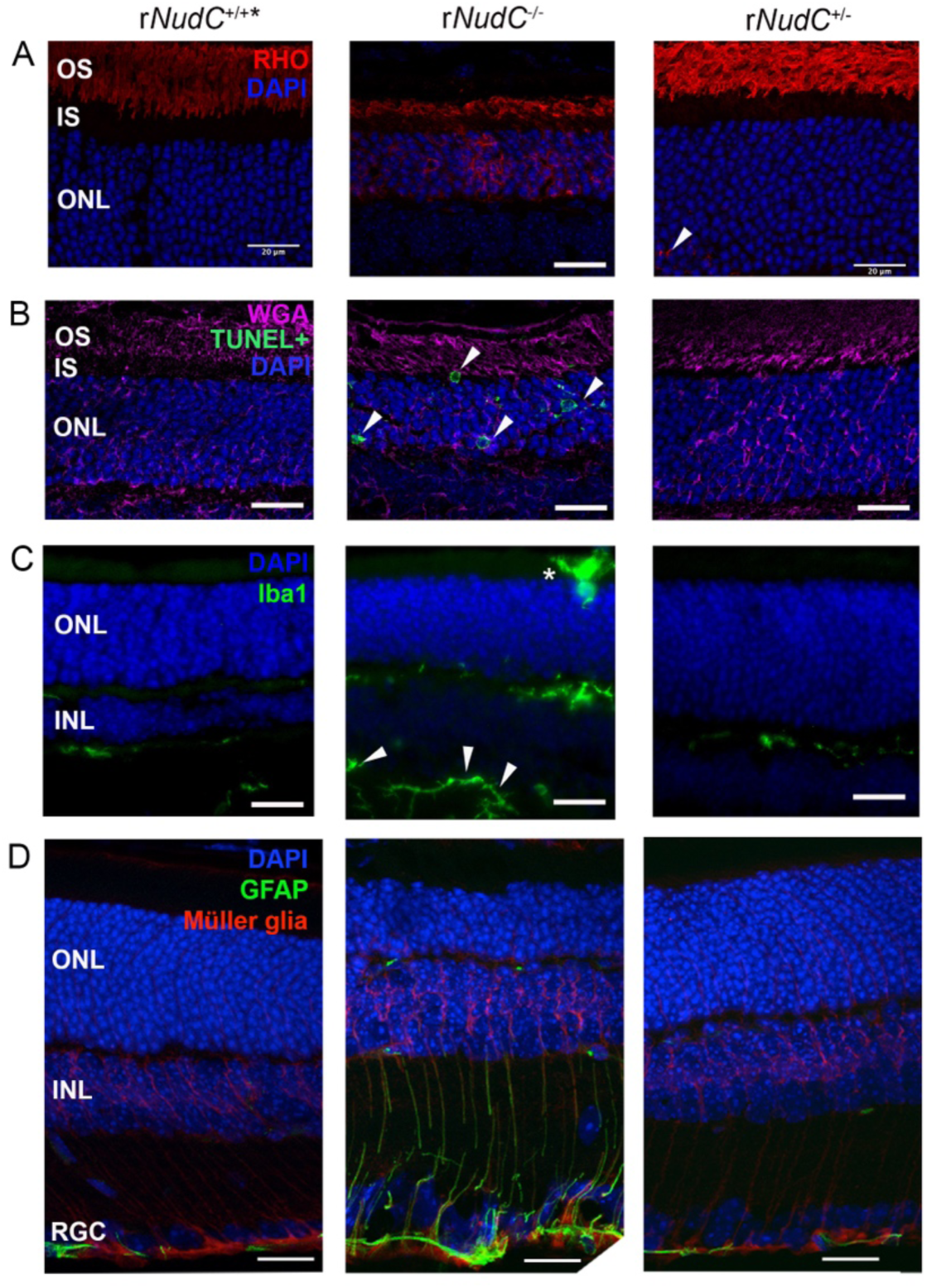
Protein mislocalization, cell death, and glial inflammatory response in 3-week-old r*NudC*^−/−^ and r*NudC*^+/−^ mouse retinas. *A*, Retinal sections stained for rhodopsin (RHO, red) displayed normal staining in the r*NudC*^+/+^ outer segment (OS) but mislocalized RHO staining in r*NudC*^−/−^ and r*NudC*^+/−^ mice (arrow) throughout the inner segment (IS) and outer nuclear layer (ONL). *B*, Retinal sections stained for apoptotic TUNEL-positive cells (green) and the OS marker, wheat germ agglutinin (WGA, pink) showed apoptotic nuclei in the ONL of r*NudC*^−/−^ (arrows) but not in r*NudC*^+/−^ or r*NudC*^+/+^ retinas. *C*, Retinal sections stained for the microglial marker Iba1 (green) showed greater microglial reactivity in the r*NudC*^-/-^ retina including in the IS (asterisk) and retinal ganglion cell layer (arrowheads). *D*, Retinal sections stained for the astrocytic marker GFAP (green) and Müller glial marker, CRALBP (red) showed increased GFAP reactivity in Müller glia in r*NudC*^-/-^ retinas. Scale bar = 20 µm. In *A-D*, nuclei are contrasted with DAPI (blue). GFAP, glial fibrillary acidic protein; INL, inner nuclear layer.

While the loss of NUDC results in a clear mislocalization of RHO at 2 (Fig. S7A) and 3 weeks (Fig. 5A), the same mice do not exhibit a mislocalization of PRPH2 by 3 weeks of age (Fig. S9).

### Loss of NUDC increases glial reactivity

Immunofluorescence revealed increased microglial reactivity via Iba1 staining in r*NudC*^-/-^ mice compared with that in r*NudC*^+/+^ mice (Fig. 5C). While microglia are normally located in the plexiform layers, as they are in r*NudC*^+/+^, in r*NudC*^-/-^, microglia were also found in the IS (Fig. 5C, asterisk) as well as in the retinal ganglion cell (RGC) layer (Fig. 5C, arrows). The r*NudC*^-/-^ mouse retina exhibited an increase in glial reactivity indicated by GFAP-positive Müller glial processes extending up through the retina (Fig. 5D, middle panel). At the 2-week timepoint, there is no evidence for Müller glial reactivity via GFAP expression (Fig. S8B).

### Cofilin 1 is found in the mouse retina in proximity to NUDC

Adult wildtype mouse retina stained for CFL1 showed staining in the IS (Fig. 6A), and proximity ligation assay for NUDC and CFL1 resulted in puncta in the IS of adult wildtype retina (Fig. 6B) indicating that NUDC and CFL1 were found within 40 nm of one another in the IS.

**Figure 6.**
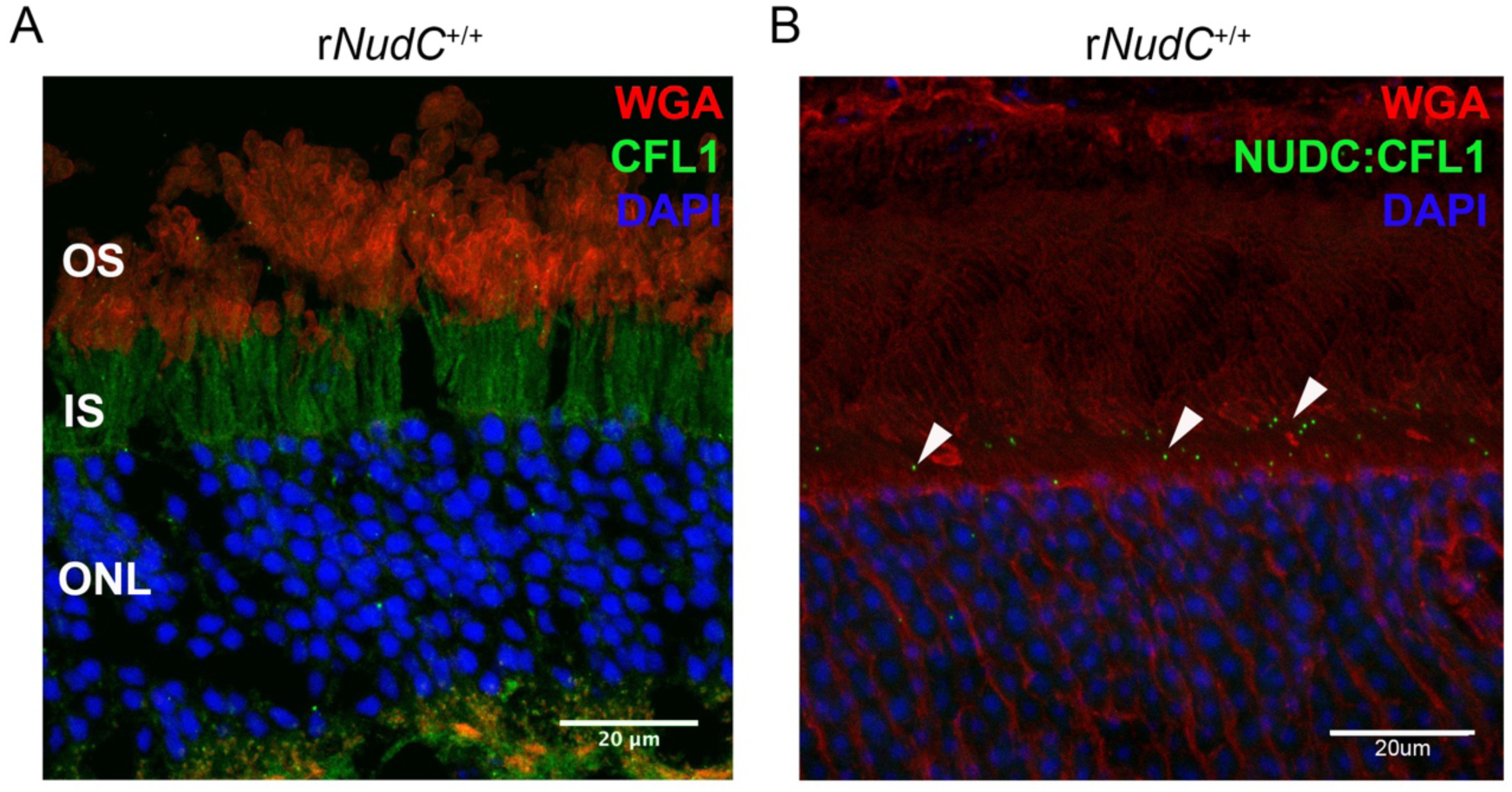
Cofilin 1 (CFL1) localization and proximity to NUDC. *A*, Adult mouse retina stained for CFL1 (green), wheat germ agglutinin (WGA, red), and the nuclear marker DAPI (blue). CFL1 is localized to the inner segment (IS). *B*, Proximity ligation assay of adult mouse retina stained for DAPI (blue), WGA (red), and CFL1: NUDC within 40 nm (green puncta) showed that CFL1 and NUDC are in close proximity to one another throughout the IS (arrows). Scale bar = 20 µm. OS, outer segment; ONL, outer nuclear layer.

### Loss of NUDC leads to actin dysregulation

When the F-actin:G-actin ratio was compared, no statistically significant differences were observed among *rNudC*^+/+^, *rNudC*^−/−^, and r*NudC*^+/−^ mice (Fig. 7A, B) at 3 weeks of age. Because NUDC interacts with the actin-remodeling protein CFL1 [30], we investigated the levels of CFL and its phosphorylated form, pCFL1, in the rod-specific *NudC* knockout. We found that although CFL1 levels remained unchanged in all genotypes, a statistically significant increase in pCFL1 was observed in *rNudC*^−/−^ mice compared with that in *rNudC*^+/+^ mice and a significant decrease was observed in the r*NudC*^+/-^ compared with r*NudC*^+/+^ (Fig. 7C, D).

**Figure 7.**
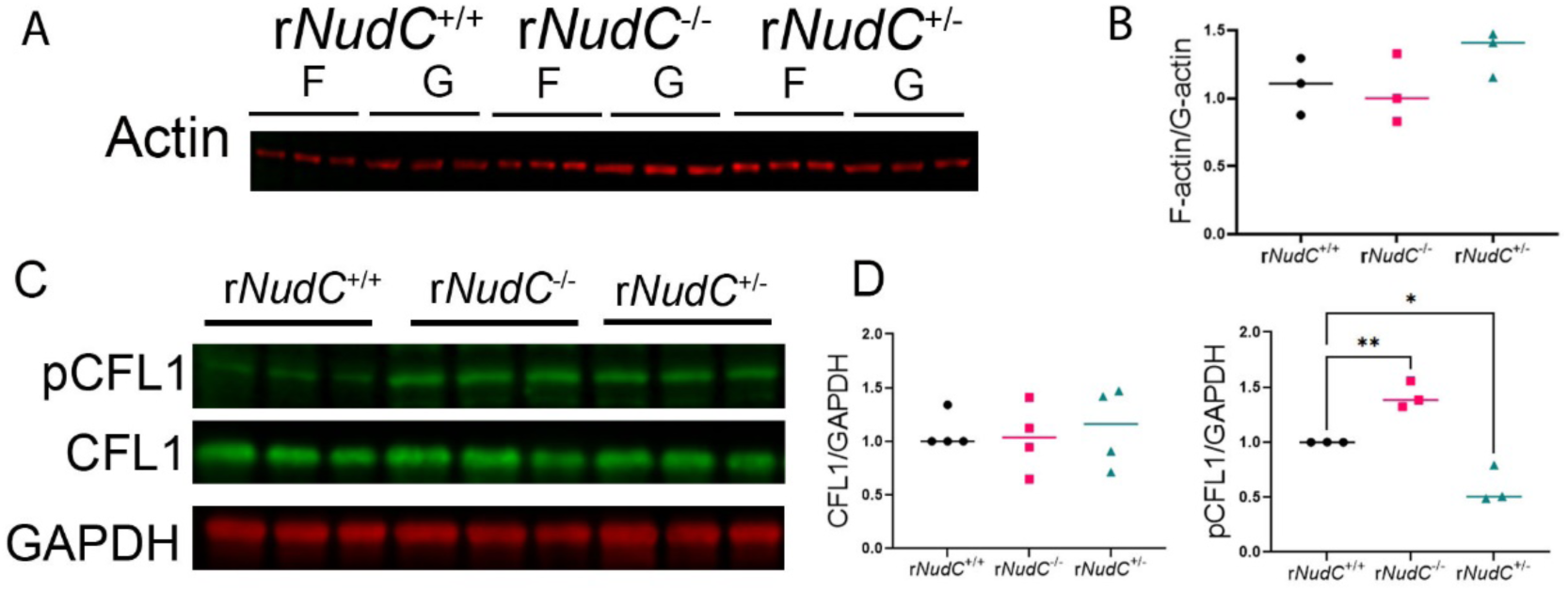
Cytoskeletal dysregulation resulting from *NudC* deletion. *A*, Western blot of F-actin and G-actin pools in retinal lysates, loaded in triplicate. *B*, Quantification of the F-actin: G-actin ratio for each genotype (n=3 for each genotype) revealed no statistically significant difference among genotypes. *C*, Western blot of retinal lysates, loaded in triplicate and probed for cofilin (CFL1) and phosphorylated CFL1 (pCFL1). *D*, Quantification of western blots in *C* revealed no change in CFL1 levels across genotypes while an increase in pCFL1 was observed in the r*NudC*^−/−^ retina compared with r*NudC*^+/+^ at 3 weeks (n=3 for each genotype). *p<0.05. F-actin, filamentous actin; G-actin, globular actin.

## Discussion

Nearly 50 years ago, proteins involved in nuclear migration were identified using the filamentous fungus, *Aspergillus nidulans*, a well-established genetic model organism for studying dynein function *in vivo* [35, 36]. Genetic analysis of the <u>nu</u>clear <u>d</u>istribution (nud) family of proteins identified a set of genes (*nudA*, *nudC*, *nudE*, *nudF*, *nudG* and *nudK*) with strong homology to mammalian dynein subunits. NudA is a homolog of the dynein heavy chain, sharing 52% identity with rat brain dynein heavy chain (DYNC1H1). NudC is 78% similar to human NUDC [37] and is required for nuclear movement [38]. In HEK293T cells, NUDC interacts with DYNLRB1 (ROBL1) and light intermediate chain 1 (DYNC1LI1 or DLIC1) [39], but not with the motor domain. NudF, a homolog of human LIS1, interacts with NudE/NDEL1 and is involved in switching dynein from the autoinhibited form to the open form to facilitate dynein activation [40, 41]. NudG is a dynein light chain of 94 amino acids with 66% identity to human dynein light chain (75% identity with human dynein light chain LC8-type 2 (DYNLL2), and *nudK* encodes ARP1 (actin-related protein 1), which is a component of the dynactin complex [1].

Deletion of *nudA* and *nudF* affect nuclear migration while deletions of the *nudC* gene in *Aspergillus* resulted in a severe phenotype, profoundly affecting the morphology and composition of the cell wall and resulting in lethality, presumably by inactivating dynein and vesicle axonemal transport [42]. Mutations in *NudC* have not been associated with human disease to date, most likely due to its well-established role in mitotic cell division where loss of NUDC would cause cell death at an early stage of fetal development. Due to this limitation, the role of NUDC as a cofactor of dynein in postmitotic neurons such as photoreceptors is incompletely understood.

In rod photoreceptors, NUDC localizes to the IS, ONL (where dynein is known to reside), OPL, and INL (**Fig. 1G**) [43]. Mouse and human *NudC* genes [37, 44] encode proteins carrying coiled-coil, nuclear localization signals (NLS), and CS domains. CS domains are protein-protein interaction domains characteristic of proteins that act as co-chaperones of Hsp90 (**Fig. S1**) and serve to stabilize proteins. To gain insight into the role of NUDC in mouse photoreceptors, we generated a rod-specific *NudC* conditional knockout facilitated by the Cre-driver iCre75 [34], which expresses Cre under control of the rhodopsin promoter. Cre expression begins at P7-8 at which point rod photoreceptors are differentiated, and the involvement of dynein in nuclei positioning and establishment of the connecting cilium [45] has already occurred. This knockout strategy allows rods to develop normally until Cre levels are high enough to delete *NudC*.

By 3 weeks of age, the NUDC knockout was evident (**Fig. 1E and F**), and by 6 weeks, r*NudC*^-/-^ rod photoreceptors were completely absent (**Fig. S4**). At 3 weeks, deletion of NUDC resulted in the loss of approximately 40% of rod photoreceptors (**Fig. 1H**) and significantly lower levels of rod OS proteins (RHO, GNAT1) (**Fig. 4**). Rhodopsin is synthesized at the rough endoplasmic reticulum surrounding rod nuclei and is normally transported via dynein to the periciliary ridge. In 2-week-old and 3-week-old r*NudC*^-/-^, rhodopsin was retained in the ONL and IS (**Figs. S8A and 5A,** middle panel, respectively) indicating that in the absence of NUDC, rhodopsin is not trafficked normally to the OS via dynein. Interestingly, peripherin-2, another protein which is localized to the OS, is not mistrafficked in the absence of NUDC (**Fig. S9**). In zebrafish neurons, loss of NUDC leads to failed dynein-cargo attachment for the initiation of retrograde transport, presumably by arresting the Phi state of dynein. This results in an accumulation of not only cargo but also dynein in the axon terminal [46]. Mitochondria, which also normally trafficked via dynein and are normally found within the IS ellipsoid region, appeared unfused and scattered throughout the *rNudC^−/−^*IS (**Fig. 2E**). As with rhodopsin mislocalization, mitochondrial mislocalization in the absence of NUDC may be due to a failure of dynein-cargo attachment.

In a rod-specific knockout of DYNC1H1, photoreceptors degenerated after P14, showing decreased scotopic ERG *a*-wave amplitudes, shortening of OS by P16, and complete photoreceptor degeneration by P30 [43]. Rod OS proteins (Rho, PDE) are reduced in the mutant OS starting at P16. In comparison, in the r*NudC*^-/-^ mice, we observed functional loss with diminished *a*- and *b*-waves at 3 weeks of age (P21) and complete loss at 6 weeks of age (P42) (**Fig. 3**). Compared to the deletion of the heavy chain, the rod degeneration of the *NudC* knockout produces a phenotype that is similar to the DYNC1H1 knockout though the rate of rod photoreceptor degeneration is slower in the absence of NUDC.

The mechanism leading to dynein inactivation by loss of NUDC is currently unclear. NUDC is required to maintain the protein levels of NUDF/LIS1 in *Aspergillus* but nudF and nudC do not co-sediment in sucrose gradients [40]. By contrast, the mammalian ortholog of NudC was identified as interacting with LIS1 in a yeast-two-hybrid screen, presumably together with other components [44], and it has been suggested that NUDC may act as a post-transcriptional regulator of NUDF/LIS1 [47]. NUDC, like LIS1, is required for neuronal migration during neocorticogenesis in radial glial progenitor cells [48] and deletions in LIS1 result in dynein inactivity and severe brain malformations, known as lissencephaly [49]. While NudC mutation leads to a near complete loss of LIS1 protein in axon terminals [46], we do not see changes in LIS1 protein levels in whole retinal lysates when NUDC is absent in rods at 3 weeks of age (**Fig. 4**). The inhibited, weakly processive state (the φ or Phi particle) of dynein [50, 51] is typically activated by binding to LIS1 and stabilizing the uninhibited open form of dynein cells [52], which enables association with dynactin and a cargo-loading effector, such as bicaudal D homolog 2 (BICD2) [53]. The fact that LIS1 levels remain unchanged at 3 weeks of age in our r*NudC*^-/-^ mouse retina could be because mouse NUDC acts upstream of LIS1 and its absence can arrest dynein in the Phi inactive form [50, 51], preventing its interaction with dynactin and cargo loading (**Fig. S10**). LIS1 may also potentially be stabilized by the NUDC-like protein 2 (NUDCL2) [54, 55].

r*NudC*^-/-^ rods showed evidence of glial reactivity (**Fig. 5C and D**) and apoptosis (**Fig. 5B**) by 3 weeks of age, and while the OSs are shorter compared to wildtype, the surviving rod OS discs appear largely structurally normal. Many remaining rods displayed an extended periciliary ridge and abnormal disc overgrowth (**Fig. 2B**, asterisks and arrowhead). This phenotype was not observed in the rod DYNC1H1 knockout but was previously recognized in a knockdown of NUDC in *X. laevis* [19]. Elongated and overgrowth of discs also occur following cytocholasin D treatment of retinal eye cups [26] and in the rod-specific knockout of *ArpC3*^-/-^ in mice P45 [56]. Recent results of photoreceptor ARPC3 (a subunit of the ARP2/3 complex) deletion demonstrated that loss of the F-actin network halted the generation of new discs and caused overgrowth of nascent discs [57]. Further, it was shown that PCARE (photoreceptor cilium actin regulator) and its downstream interactor, WASF3 (Wiskott-Aldrich syndrome protein family member 3) regulate ciliary F-actin assembly and ciliary membrane expansion during disc morphogenesis [58].

Our results show loss of NUDC in rods affected actin cytoskeleton regulation. NUDC is known to stabilize CFL1, which is a potent regulator of actin filament dynamics. The ability of CFL1 to bind and depolymerize actin is abolished by phosphorylation of serine residue 4 by LIMK1 [59, 60]. We found CFL1 to be localized within close proximity (40 nm) to NUDC in the rod IS (**Fig. 6B**), and loss of NUDC resulted in an increase in the phosphorylated form of CFL1 (pCFL1). This increase in pCFL1 indicates that a significant amount of CFL1 was no longer bound to F-actin, and a portion of actin was no longer being clipped to form G-actin or branched F-actin. We expected to see an increase in the F-actin: G-actin ratio as a result of the loss of NUDC; however, r*NudC*^-/-^ mice at 3 weeks of age did not reveal a difference compared with r*NudC*^+/+^ (**Fig. 7**). This may be because a 40% loss of rods was already observed at this stage in r*NudC*^−/−^ mice, and any change in actin pools would not be evident in the remaining rods because we measured the total F-actin: G-actin ratio in whole retinal lysates. There was, however, a slight (though not significant) increase in F-actin: G-actin in r*NudC*^+/−^ mice at 3 weeks as well as a statistically significant decrease in pCFL.

While well-established as necessary for cell division, our results indicate a role for NUDC in dynein-dependent protein and organelle trafficking as well as in actin cytoskeletal dynamics in postmitotic rod photoreceptors (**Fig. 8**). Deletion of NUDC in rods presumably arrests dynein in the inactive Phi form, affecting protein and organelle trafficking in postmitotic rods, and loss of NUDC also affects pCFL1 levels and therefore actin cytoskeletal dynamics. These findings establish a critical role for NUDC in postmitotic rod photoreceptor maintenance and survival.

**Figure 8.**
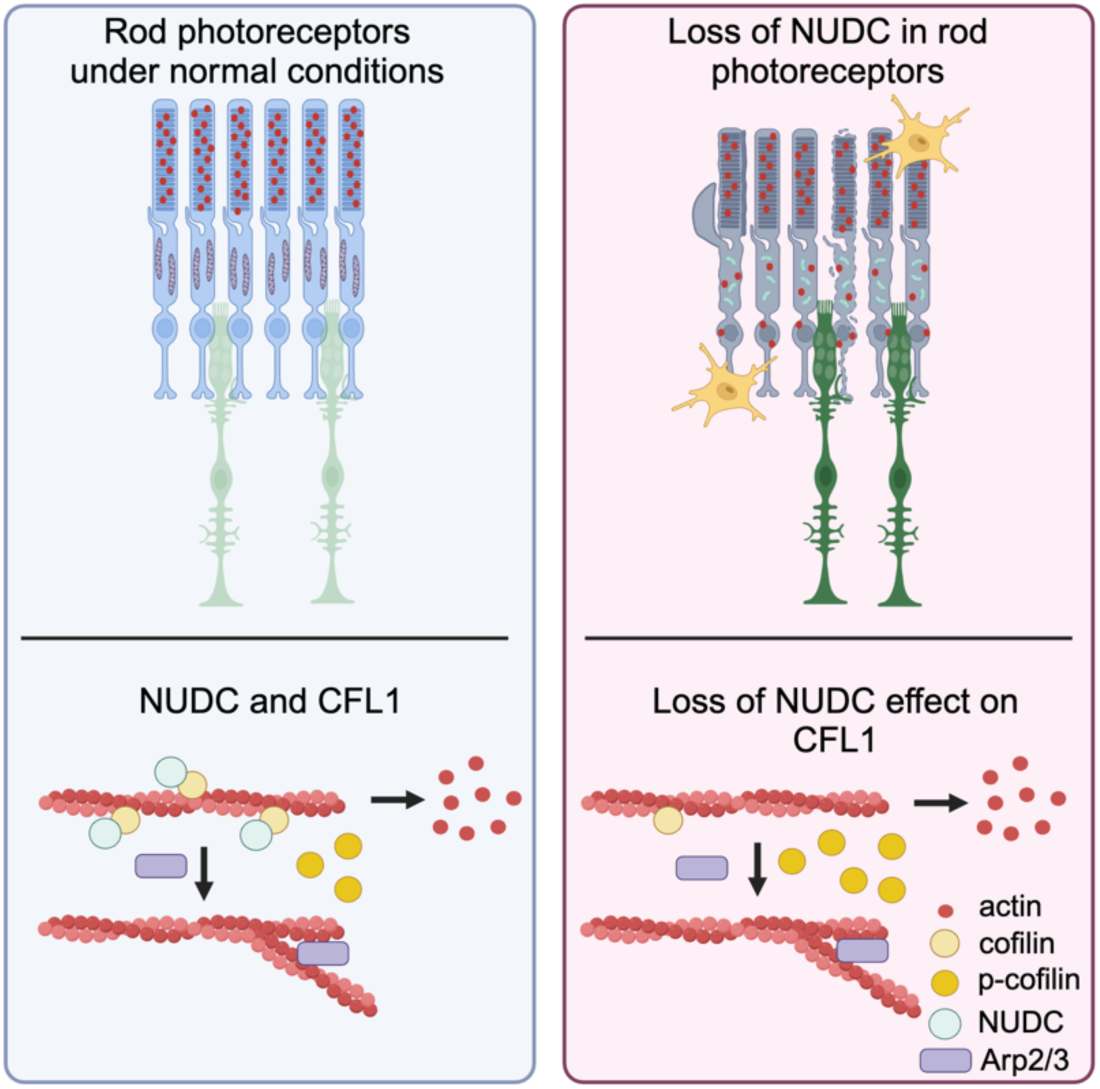
Schematic diagram of mouse rod photoreceptors under normal conditions (left panel) or in the absence of NUDC in rods (right panel) at 3 weeks of age. In r*NudC*^+/+^ mouse rods (left), rhodopsin (RHO, red circles) is properly trafficked to the outer segment (OS), and mitochondria are elongated and located next to the plasma membrane. In *rNudC*^−/−^ mouse rods (right), the mitochondria appear smaller in ultrathin sections and scattered throughout the inner segment (IS), and some rods exhibit an extended periciliary ridge. Additionally, RHO is mislocalized in the IS and outer nuclear layer (ONL) of the retina. Müller glia (green cells) express GFAP throughout their processes, and microglia (yellow cells) migrate towards the outer retina. While NUDC normally stabilizes CFL1, in the absence of NUDC, there is an increase in the proportion of CFL1 that is phosphorylated (pCFL1).

## Materials and Methods

### Research animals

Flippase (FLP) and iCre75 transgenic mice were obtained from The Jackson Laboratory (Bar Harbor, ME, USA). Mice were maintained under 12-hour cyclic dark/light conditions in standard cages and fed *ad libitum*. All animal studies were conducted in compliance with the National Institutes of Health *Guide for the Care and Use of Laboratory Animals* and were approved by the Institutional Animal Care and Use Committee of the University of Alabama at Birmingham.

### Generation of rod-specific *NudC*^−/−^ mice

NudC embryonic stem (ES) cells were purchased from the KOMP (Knockout Mouse Project) and developed at the University of Utah Transgenic Gene-Targeting Mouse Facility. The construct of this clone contained a gene trap (GT) cassette inserted into intron 2 and floxed exons 3 and 7 (see Fig. 1A for diagram). Gene targeting of ES cells was confirmed using PCR, including the 3’ and 5’ recombination arm, FRT sites, and LoxP sites. ES cell blastocyst injection and generation of chimeric and heterozygous *NudC*^tm1a^ (GT/+) mice was performed at the University of Utah transgenic core. Homozygous gene-trapped mice did not show a phenotype indicating that the gene did not block translation of NUDC (“leaky” gene trap). Next, we crossed NudC^GT/+^ with FLP (ROSA26::FLPe) knockin mice (C57BL/6 background) to yield animals with the *NudC*^tm1c^ floxed allele (NudC^fl/+^). The presence of the floxed *NudC* allele was confirmed using PCR with the following primers flanking the third loxP site: a) 5’ GTAAGCAACTTGGGCATCAG 3’ and b) 5’ CTAGCCAAGACAGAACCACG 3’. Homozygous *NudC*^fl/fl^ mice were crossed with rod photoreceptor-specific Cre recombinase (iCre75) mice to generate *NudC*^tm1d^ rod-specific r*NudC*^+/−^ and r*NudC*^−/−^ mice. iCre75 mice expressed Cre recombinase specifically in rod photoreceptors from P4–P7 with total excision completed by P18. The presence of iCre75 was confirmed using PCR with the primers 5’ GGATGCCACCTCTGATGAAG 3’ and 5’ CACACCATTCTTTCTGACCG 3’, and animals were bred such that there was only one iCre75-expressing allele. Excision of exons 3–7 was confirmed by PCR of retina lysates using the following primers: c) 5’ GCAGTTTCTGAGTCTGTTGAGAG 3’, d) 5’ TCCTGATCCTCTGCATCCTTTT 3’, and e) 5’ GCAGAAGTGGAAAAGTGACCTC.

### Electroretinography

ERG was performed on 3-week-old and 6-week-old wild-type (r*NudC*^+/+^), heterozygous (r*NudC*^+/−^), knockout (r*NudC*^−/−^), and control (r*NudC*^fl/fl^iCre75^-^ and r*NudC*^+/+^iCre75^+^) mice (n=3–4 each group) using an LKC UTAS-3000 Visual Electrodiagnostic System (Gaithersburg, MD, USA) as described previously [61]. Briefly, mice were dark-adapted overnight and anesthetized by intraperitoneal injection of ketamine (71.5 μg/g) and xylazine (14.3 μg/g) in 0.1M phosphate-buffered saline (PBS), and the pupils were dilated with a mixture of 0.5% phenylephrine, 0.5% proparacaine HCl, and 1% tropicamide. Silver-embedded thread electrodes were placed in contact with the corneal surface, which was covered with 2.5% hypromellose ophthalmic solution (GONIOVISC). Scotopic ERGs were performed at the following light intensities: −20 dB (0.025 cd*s/m^2^), −10 dB (0.25 cd*s/m^2^), 0 dB (2.5 cd*s/m^2^), 5 dB (7.91 cd*s/m^2^), 10 dB (25 cd*s/m^2^), and 15 dB (79.1 cd*s/m^2^). Five scans were performed and averaged for each light intensity. The *a*-wave amplitudes were measured from the baseline to trough, and the *b*-wave amplitudes were determined from baseline to the major cornea-positive peak. Photopic ERGs were measured after a 10-minute exposure to 25 cd*s/m^2^ background light followed by a range of flash intensities. Latencies represent the time (ms) from the peak or trough. Data were collected using Microsoft Excel and graphed using GraphPad Prism.

### Transmission electron microscopy

Following euthanasia, retinal tissues were harvested and fixed in mixed aldehyde solutions (2% paraformaldehyde, 2% glutaraldehyde, 0.05% calcium chloride in 50 mM MOPS, pH 7.4), then osmicated, dehydrated, resin embedded in Epon and sectioned at 70 nm. Sections were placed on copper grids and stained with 2% uranyl acetate and 3.5% lead citrate (19314; Ted Pella). Images were acquired at 2.18 nm/px with a LEOL JEM-1400 TEM equipped with a 16-Mpixel Gatan camera using SerialEM software. Image mosaics were assembled with Nornir software application. Images were also acquired on a Tecnai Spirit T12 Transmission Electron Microscope (Thermo Fisher, formerly FEI) with an AMT (Advance Microscopy Techniques, Corp) Bio Sprint 29-megapixel digital camera.

### Immunofluorescence

Enucleated eyes from 3-week-old mice of either sex were placed in 4% paraformaldehyde in 0.1 M PBS (pH 7.4) overnight at 4°C. Following an additional overnight incubation in 30% sucrose in PBS at 4°C, the eyes were embedded in Tissue-Tek O.C.T. (Sakura Finetek, Torrance, CA, USA), flash frozen in isopentane cooled to near freezing by liquid nitrogen and sectioned into 10-μm-thick slices using a cryomicrotome. Following sectioning, the eyes were processed for IHC as previously described [19]. The sections were incubated in 10 mM sodium citrate and 0.05% Tween (pH 6.0) for 1 hour for heat-induced epitope retrieval and cooled to room temperature for 30 minutes. Primary antibodies (anti-NUDC, Abcam, 1:400; anti-GFAP, Cell Signaling, 1:400; anti-CRALBP (Müller glia marker), Abcam, 1:250; anti-Iba1, Cell Signaling, 1:100; anti-rhodopsin B6-30N, a gift from Drs. Paul Hargrave and W. Clay Smith (University of Florida), 1:500; and wheat germ agglutinin (1:750) Thermo Fisher Scientific, Waltham, MA, USA, conjugated with Alexa Fluor 555 or Alexa Fluor 647) were used to counterstain the photoreceptor OS. For TUNEL labeling, the Click-It Plus TUNEL kit (Thermo Fisher Scientific) was used to label apoptotic nuclei according to the manufacturer’s instructions. Images of stained retinal sections were acquired on a Zeiss LSM 800 Airyscan confocal microscope (Carl Zeiss, Oberkochen, Germany) with a 63× oil-immersion objective attached to an Axiocam-506 CCD camera (Carl Zeiss) or using an ECHO Revolution (ECHO, BIO Convergence Company). ImageJ software (NIH) was used to compile maximum projection *z*-stacks and process acquired images.

### Proximity Ligation Assay

Cryosections (10mm) or fixed retinal tissue were incubated in 10 mM sodium citrate and 0.05% Tween (pH 6.0) for 1 hour for heat-induced epitope retrieval and cooled to room temperature for 30 minutes prior to following the protocol for Duolink PLA Fluorescence (Sigma). Briefly, slides were blocked in Duolink blocking solution for 1 hour at 37°C in a humidified chamber prior to primary antibody incubation overnight at 4°C in a humidified chamber. Primary antibodies used include anti-NUDC, Abcam, 1:400; anti-NUDC, Santa Cruz, 1:100; anti-CFL1, Abcam, 5 μg/ml; anti-CFL1, Cell Signaling, 1:400. Following two washes in Wash Buffer A, (WBA) retinal slices were incubated with Duolink Probes diluted 1:5 for 1 hour at 37°C. Retinal slices were incubated in ligation buffer, washed twice with WBA, and exposed to ligase for 30 minutes at 37°C. Amplification in the presence of polymerase for 100 minutes at 37°C was followed by a series of washes in Wash Buffer B prior to mounting with Duolink Mounting Medium with DAPI. Slides were then stored at 4°C until imaging using a Zeiss LSM 800 Airyscan confocal microscope (Carl Zeiss, Oberkochen, Germany).

### Spidergrams

Cryosections (10 μm) of fixed retinal tissue were stained with hematoxylin and eosin as previously described [62], and brightfield images were obtained using an ECHO Revolution (ECHO, BIO Convergence Company) at 20× and joined together to produce an image of the entire retina, including the optic nerve. ImageJ software was used to measure the distance from the optic nerve and count nuclei in each 200-μm segment of the ONL.

### Western blotting

Mouse retinas from both eyes of each genotype, with n=3 separate animals per genotype, were homogenized in 100 μl lysis buffer (10 mM Tris, Ph 7.2, 5 mM EDTA, 150 mM NaCl, 1% Triton x100, 0.5% NP40 plus protease and phosphatase inhibitor cocktails), and protein concentrations were quantified using a BCA protein assay (Pierce). Precast 12% SDS Page Criterion gels (BioRad) were loaded based on the linear range of each antibody (20 μg for each antibody except for PHPR2, which was loaded with 10 μg to remain within the linear range), and each sample was run in triplicate at 200 V for 45 minutes. The proteins were transferred onto nitrocellulose membranes using a semi-dry transfer system (BioRad). The membranes were blocked in 5% milk in TBS-T before incubation with primary antibodies overnight at 4°C in 3% milk or BSA in TBS-T. The primary antibodies used included anti-NUDC (Abcam, 1:500); anti-arrestin (Cell Signaling Technologies [CST], 1:500); anti-transducin (Abcam, 1:2000); anti-LIS1 (CST, 1:1μg/ml); anti-Rab11A (Abcam, 1 μg/mL); anti-rhodopsin (anti-rhodopsin 1D4, a gift from Dr. Robert Molday (University of British Columbia), 1 mg/mL); anti-peripherin (EMD, 1:1000); anti-CFL1 (CST, 1:1000); anti-pCFL1 (CST, 1:500); and the loading control, anti-GAPDH (CST, 1:1000 or Abcam, 1:10,000). AlexaFluor anti-rabbit or mouse secondary antibodies (1:10,000) were incubated in 3% milk or BSA in TBS-T for 1 hour at room temperature, and the membranes were visualized using a Licor Odyssey Fc NIR imaging system (LI-COR, Lincoln, NE, USA). Bands were quantified using ImageJ software, and GAPDH was used as a loading control.

Western blots to determine the ratio of G-actin to F-actin were performed on retinal lysates that were processed using a G-actin/F-actin In Vivo Assay Kit (Cytoskeleton, Inc.) following the manufacturer’s instructions. Briefly, retinal tissue from both eyes of each genotype, with n=3 separate animals per genotype, were homogenized in 100 μl of 37°C LAS2 and incubated at 37°C for 10 minutes before centrifugation at 350×g for 5 minutes at room temperature to pellet debris. The supernatant was removed and placed in a fresh tube before centrifugation at 100,000×g at 37°C for 1 hour. Following centrifugation, the supernatant (G-actin pool) was placed in a fresh tube, and F-actin depolymerization buffer was added to the pellet (F-actin pool) and incubated on ice for 1 hour, pipetting every 15 minutes. SDS sample buffer (5×) was added to the pellet and supernatant fractions and mixed well. The samples were stored at −20°C prior to western blot analysis for actin (Cytoskeleton 1:1000). Standards (10, 20, and 50 ng) were resuspended in F-actin depolymerization buffer, and 12% SDS Page Criterion gels (BioRad) were loaded 10 μl per sample prior to being run as described above. Primary antibody, anti-α-actin (Cytoskeleton, Inc., 1:1000) was incubated overnight at 4°C in 3% BSA in TBS-T. AlexaFluor anti-mouse antibody (1:10,000) was incubated in 3% BSA in TBS-T for 1 hour at room temperature, and membranes were visualized as described above. Unlike a typical western blot, this assay results in a single sample being separated into fractions to produce pools of F- and G-actin and therefore does not utilize a loading control such as GAPDH.

### Statistical analysis

Data are presented as the mean±standard deviation, where n represents the number of mice analyzed. Statistical comparisons were performed using one-way ANOVA for all experimental data. A value of p<0.05 indicates a statistically significant difference.

## Author Contributions

Mary Anne Garner wrote the first draft and subsequent edits of the manuscript, and performed experiments (Figs. 1E-H, 4, 5C and D, 7, 8, S2, S3, S8, S9); Meredith G. Hubbard performed experiments (Figs. 2, 3, S4, S5, S6, S7); Evan R. Boitet performed experiments (Figs. 5A–B, 6); Seth T. Hubbard performed experiments (Figs. 1H, S2); Anushree Gade performed experiments (Fig. 1D); Guoxin Ying helped create the knockout mice (Fig. S1); Bryan W. Jones performed experiments (Fig. 2, S3); Wolfgang Baehr helped create the knockout mice (Fig. S1), developed the model interaction of NUDC with dynein (Fig. S10) and edited the manuscript; Alecia K. Gross conceptualized the project, helped create the knockout mice (Fig. S1), completed the experimental design, and edited the manuscript.

## Data Availability Statement

The authors confirm that the data supporting the findings of this study are available within the article and its supplementary materials and are also available from the corresponding author, AKG, upon reasonable request.

## Acknowledgments

We would like to express our gratitude to Drs. Paul Hargrave and Clay Smith for the use of their B6-30N Rhodopsin antibody and to Dr. Robert Molday for the use of the 1D4 Rhodopsin antibody. We acknowledge assistance from Melissa Chimento in the UAB High Resolution Imaging Facility. We would also like to thank Mr. Jean Sun for offering technical advice during the preparation of this manuscript. This research was supported by the National Eye Institute (NEI) grants EY030096, EY019311 (AKG); EY019298 (WB); EY028927 (BWJ); EY014800-039003 (Moran Eye Center); P30 EY003039 (UAB); NSF 2014862 (BWJ), an unrestricted grant from Gabe Newell to BWJ, an unrestricted grant to the Department of Ophthalmology and Visual Sciences, University of Utah from Research to Prevent Blindness (RPB; New York); grant from the Retina Research Foundation-Houston to WB (Alice McPherson, MD).

