## Supplemental Figures 1-10 for "NUDC is critical for rod photoreceptor function, maintenance, and survival"


**This PDF file includes:**

Figures S1 to S10

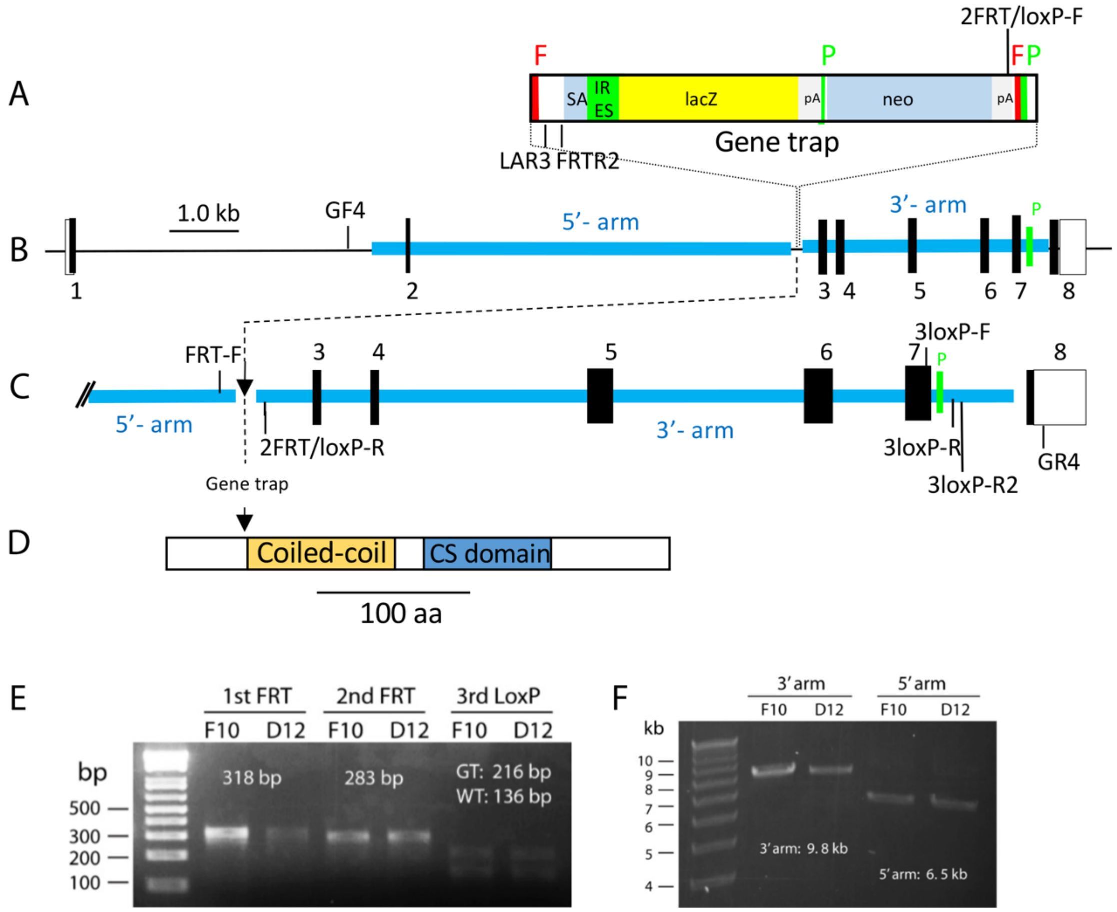


**Fig. S1.** NUDC knockout strategy using *Nudc* tm1a(KOMP)Mbp stem cell line. *A*, gene trap with lacZ and neo cassettes positioned in intron 2. SA, splice acceptor; IRES, internal ribosome entry site; pA, polyadenylation site; F, FRT; P, loxP. *B*, schematic of *Nudc* gene with 8 exons. Blue lines, 5'- and 3'-arms used for homologous recombination. *C*, enlargement of the *NudC* gene. Gene trap was inserted between 5'-arm and 3'-arm. GF4, GR4, loxP-F/R, LAR3 denote primers. *D*, schematic of mouse NUDC with functional domain coiled-coil and CS. Arrow indicates truncation point in NUDC after exon 1 and K-53. *E*, verification of first FRT, second FRT and third loxP in *NudC* tm1a by PCR. F10 and D12 denote two different embryonic cell line clones. F10 was used to generate the floxed allele. *F*, verification of 5'-arm (long arm) and 3'-arm (short arm) in *NudC* tm1a F10 and D12.

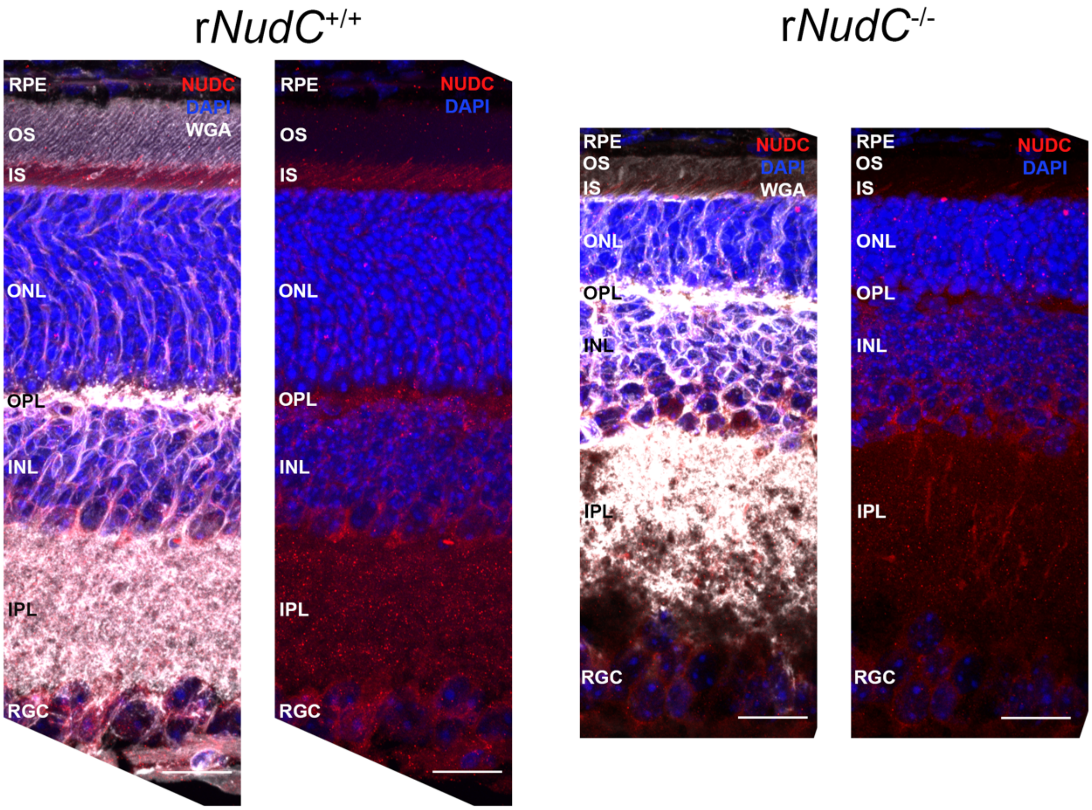


**Fig. S2.** NUDC expression across the entire mouse retina at 3 weeks of age. *A*, Retinal sections stained for NUDC (red), wheat germ agglutinin (WGA, grey), and the nuclear marker DAPI (blue) displayed normal staining in the r*NudC*$^{+/+}$ retinal pigmented epithelium (RPE), inner segment (IS), outer nuclear layer (ONL), outer plexiform layer (OPL), inner nuclear layer (INL), inner plexiform layer (IPL), and retinal ganglion cell layer (RGC). *B*, NUDC staining in r*NudC*$^{-/-}$ is diminished in the IS and ONL but is still present in the RPE, OPL, INL, IPL, and RGC layers. Scalebar = 20 μm.

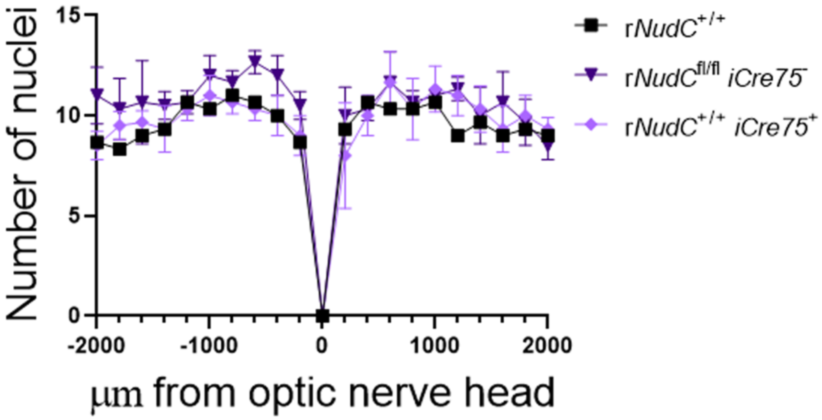


**Fig. S3.** Spidergram of control mice at 3 weeks of age. Quantification of outer nuclear layer (ONL) nuclei counts every 200 µm from the optic nerve head, n=3 for each genotype. OS, outer segment; IS, inner segment. Scale bar = 20 µm.

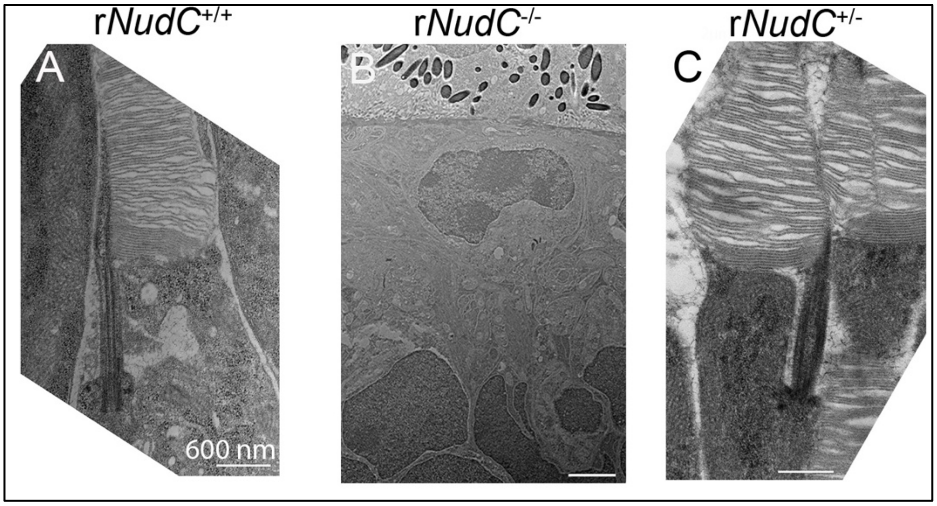


**Fig. S4.** Transmission electron microscopy (TEM). A-C, images taken from 6-week-old mouse photoreceptors. A-C, scale bar = 600 nm. Absence of NUDC causes complete outer segment degeneration by 6 weeks.

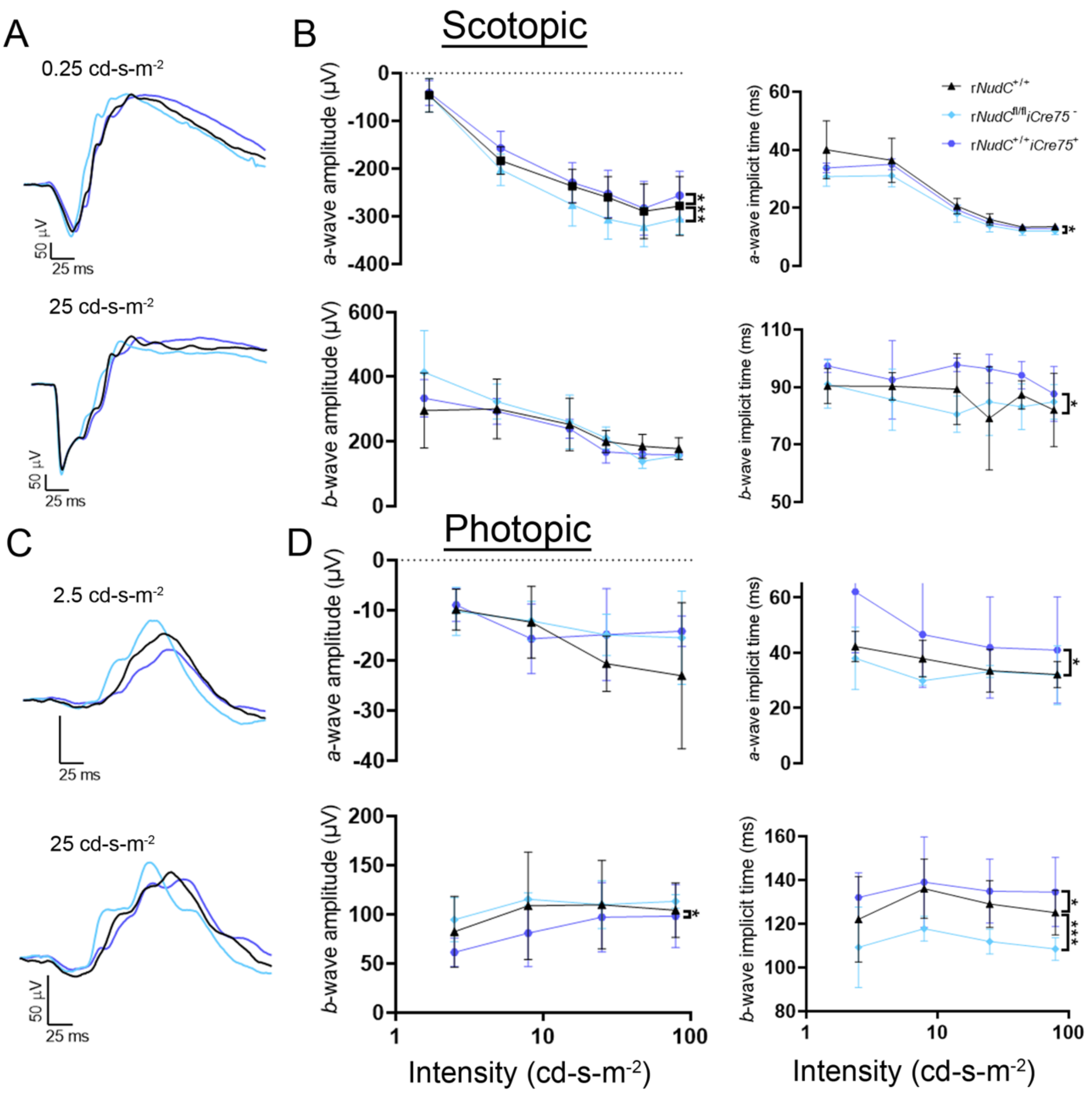


**Fig. S5.** Retinal function in 3-week-old rod-specific *NudC* control mice. *A*, Average ERG (n=3) from the lowest (0.25 cd s/m$^2$) and highest (25 cd s/m$^2$) flash intensities under scotopic conditions for 3-week-old mice. r*NudC*$^{+/+}$, black traces; r*NudC*$^{fl/fl}$*Cre75*$^{-}$, light blue traces; r*NudC*$^{+/+}$ *iCre75*$^{-}$, dark blue traces. *B*, Summary data of *a*-wave and *b*-wave amplitudes and implicit times as a function of light intensity from 3-week-old mice. r*NudC*$^{+/+}$, black; r*NudC*$^{fl/fl}$ *iCre75*$^{-}$, light blue; r*NudC*$^{+/+}$ *iCre75*$^{-}$, dark blue. *p<0.05, ***p<0.0001. *C*, Average ERG recordings (n=3) from the lowest (2.5 cd s/m$^2$) and highest (25 cd s/m$^2$) flash intensities under photopic conditions for 3-week-old mice. r*NudC*$^{+/+}$, black traces; r*NudC*$^{fl/fl}$ *iCre75*$^{-}$, light blue traces; r*NudC*$^{+/+}$ *iCre75*$^{-}$, dark blue traces. *D*, Summary data of *a*-wave and *b*-wave amplitudes and implicit times as a function of light intensity from 3-week-old mice. r*NudC*$^{+/+}$, black; r*NudC*$^{fl/fl}$ *iCre75*$^{-}$, light blue; r*NudC*$^{+/+}$ *iCre75*$^{-}$, dark blue. *p<0.05, ***p<0.0001.

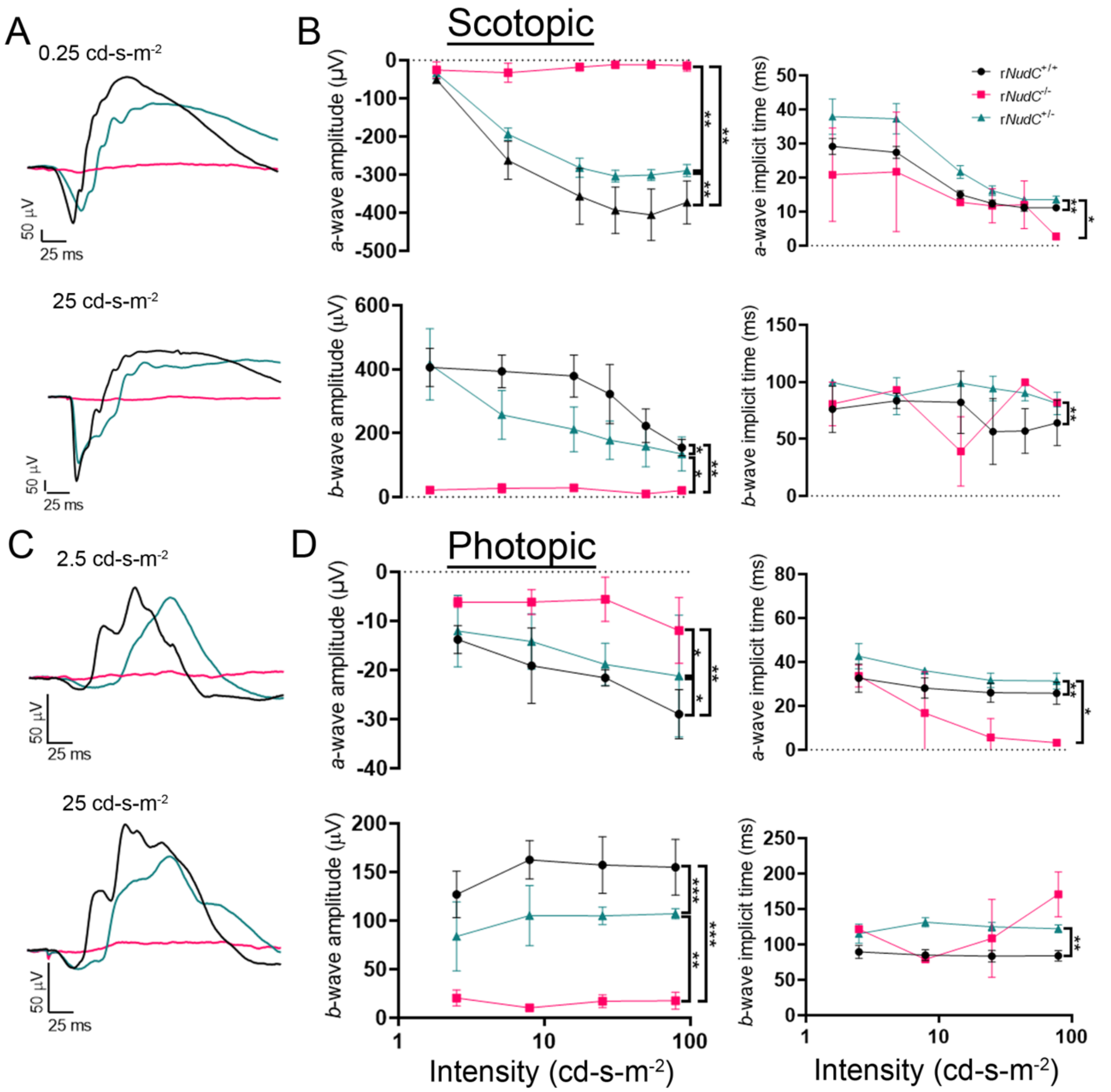


**Fig. S6.** Loss of *NudC* affects retinal function in 6-week-old mice. *A*, Average electroretinography recordings (n=3) from the lowest (0.25 cd s/$m^2$) and highest (25 cd s/$m^2$) flash intensities under scotopic conditions. r*NudC*$^{+/+}$, black traces; r*NudC*$^{-/-}$, pink traces; r*NudC*$^{+/-}$, green traces. *B*, Summary data of *a*-wave and *b*-wave amplitudes and implicit times as a function of light intensity. r*NudC*$^{+/+}$, black; r*NudC*$^{-/-}$, pink; r*NudC*$^{+/-}$, green. *p<0.05, **p<0.001, ***p<0.0001. *C*, Average ERG recordings (n=3) from the lowest (2.5 cd s/$m^2$) and highest (25 cd s/$m^2$) flash intensities under photopic conditions. r*NudC*$^{+/+}$, black traces; r*NudC*$^{-/-}$, pink traces; r*NudC*$^{+/-}$, green traces. *D*, Summary data of *a*-wave and *b*-wave amplitudes and implicit times as a function of light intensity. r*NudC*$^{+/+}$, black; r*NudC*$^{-/-}$, pink; r*NudC*$^{+/-}$, green. * p<0.05, **p< 0.001, ***p<0.0001.

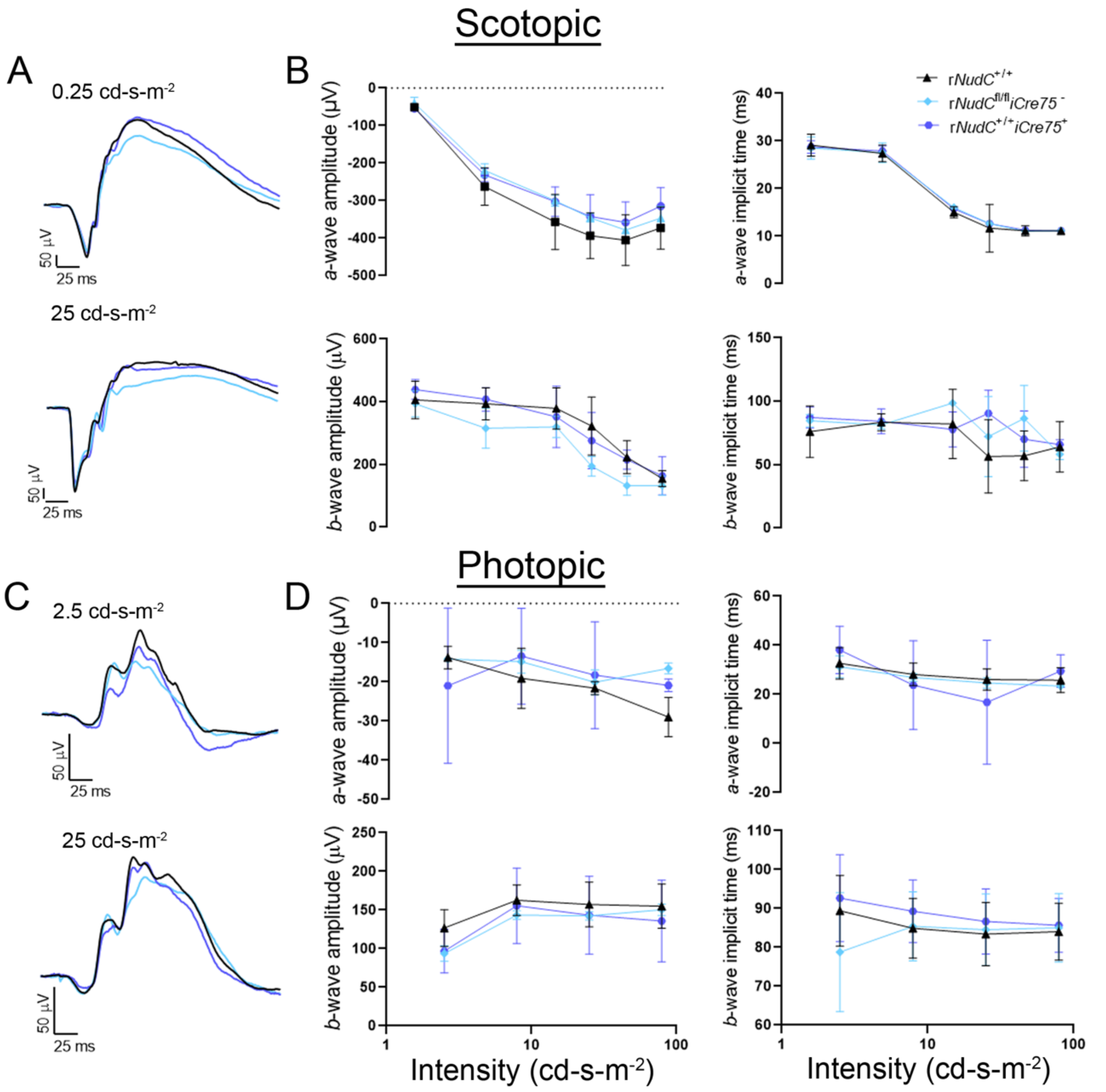


**Fig. S7.** Retinal function in 6-week-old rod-specific *NudC* control mice. *A*, Average electroretinography recordings (n=3) from the lowest (0.25 cd s/m$^2$) and highest (25 cd s/m$^2$) flash intensities under scotopic conditions. r*NudC*$^{+/+}$, black traces; r*NudC*$^{fl/fl}$ *iCre75*$^-$, light blue traces; r*NudC*$^{+/+}$ *iCre75*$^-$, dark blue traces. *B*, Summary data of *a*-wave and *b*-wave amplitudes and implicit times as a function of light intensity. r*NudC*$^{+/+}$, black; r*NudC*$^{fl/fl}$ *iCre75*$^-$, light blue; r*NudC*$^{+/+}$ *iCre75*$^-$, dark blue. *C*, Average electroretinography recordings (n=3) from the lowest (2.5 cd s/m$^2$) and highest (25 cd s/m$^2$) flash intensities under photopic conditions. r*NudC*$^{+/+}$, black traces; r*NudC*$^{fl/fl}$ *iCre75*$^-$, light blue traces; r*NudC*$^{+/+}$ *iCre75*$^-$, dark blue traces. *D*, Summary data of *a*-wave and *b*-wave amplitudes and implicit times as a function of light intensity. r*NudC*$^{+/+}$, black; r*NudC*$^{fl/fl}$ *iCre75*$^-$, light blue; r*NudC*$^{+/+}$ *iCre75*$^-$, dark blue.

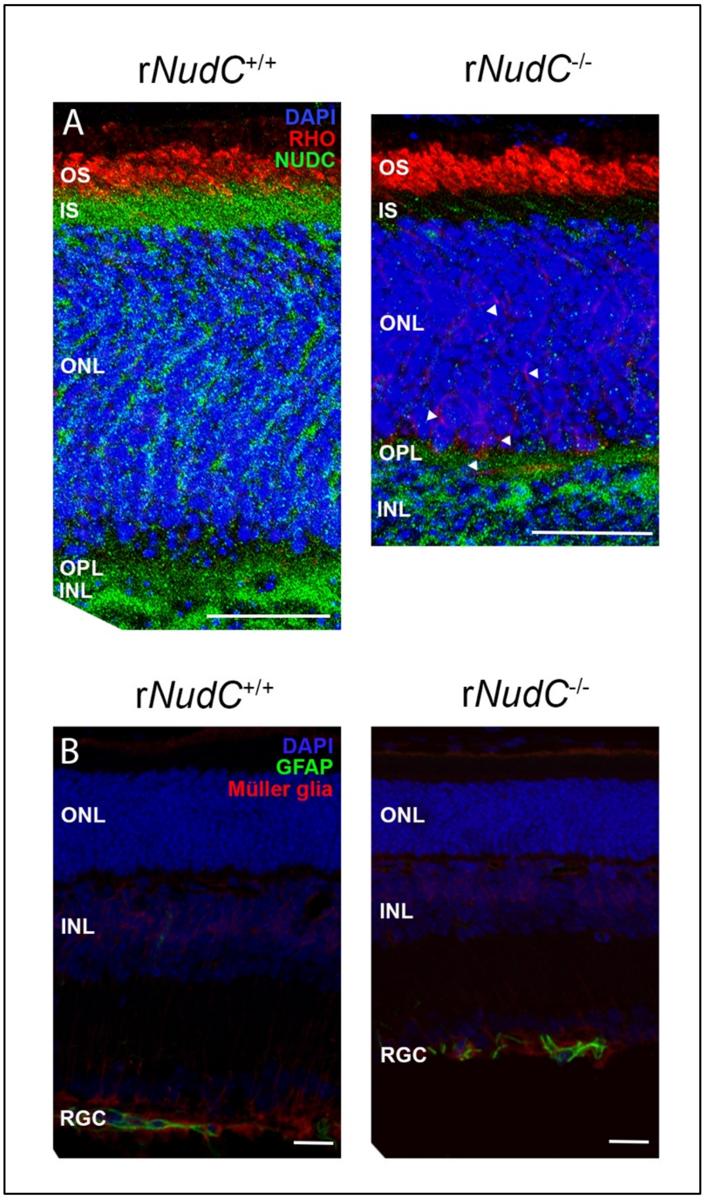


**Fig. S8.** NUDC loss, RHO mislocalization, and lack of Müller glia reactivity in 2-week-old r*NudC*$^{-/-}$ mouse retina. *A*, Retinal sections stained for rhodopsin (RHO, red) displayed normal staining in the r*NudC*$^{+/+}$ outer segment (OS) but mislocalized RHO staining in r*NudC*$^{-/-}$ (arrowheads) throughout the outer nuclear layer (ONL). *B*, Retinal sections stained for the astrocytic marker GFAP (green) and Müller glial marker, CRALBP (red) showed no difference in GFAP reactivity in in 2-week-old r*NudC*$^{-/-}$ retinas. Scalebar = 20 µm.

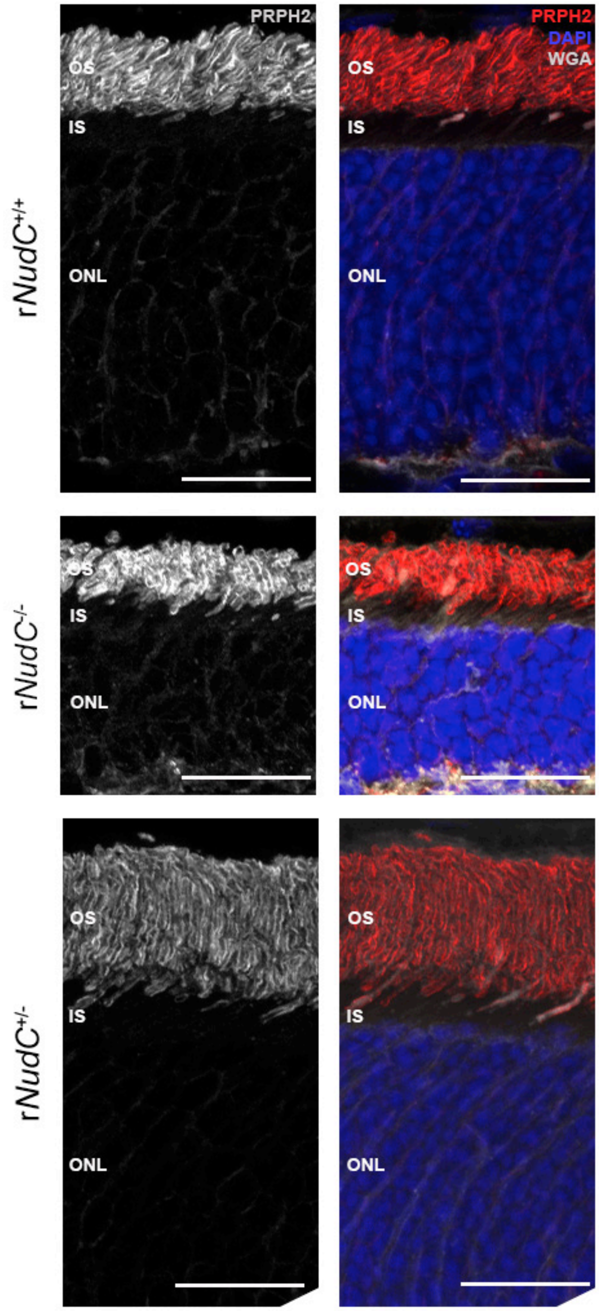


**Fig. S9.** Peripherin localization in 3-week-old r*NudC*$^{-/-}$ and r*NudC*$^{+/-}$ mouse retinas. 3-week-old mice were stained for peripherin-2 (PRPH2, red), wheat germ agglutinin (WGA, grey), and the nuclear marker DAPI (blue). Peripherin is properly localized to the outer segment (OS) in the r*NudC*$^{+/+}$, r*NudC*$^{-/-}$, and r*NudC*$^{+/-}$retinas. Scale bar = 20 μm. OS, outer segment; IS, inner segment; ONL, outer nuclear layer.

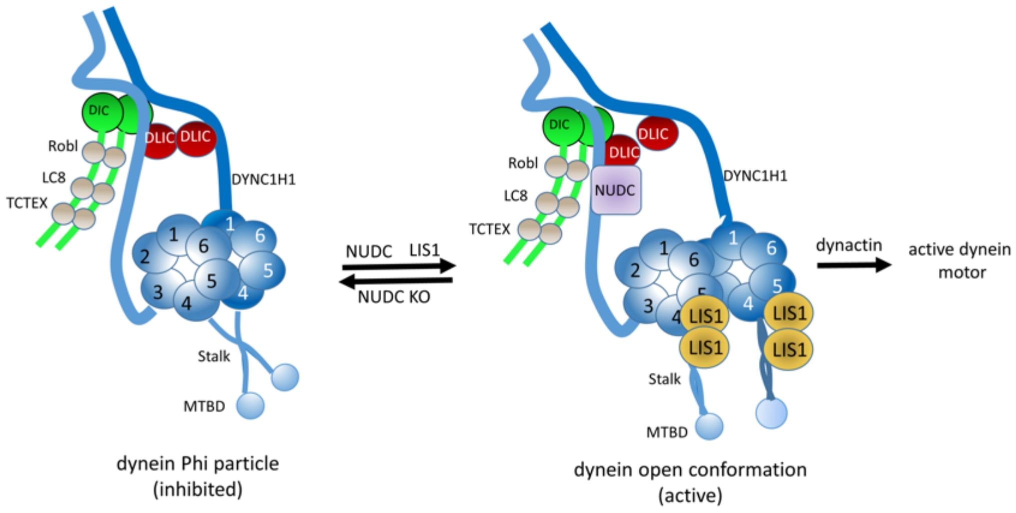


**Fig. S10.** Schematic of dynein activation. *A*, representation of multimeric dynein in the inhibited Phi form. The heavy chains (DYNC1H1) form homodimers, the scaffolds of which organize the distributions of heterodimeric intermediate (DIC), light intermediate (DLIC), and light chains (ROBL, LC8, TCTEX) (adapted from Dahl, 2021). Blue circles 1-6 represent the motor domain. *B*, active dynein in the open conformation. NUDC interacts with dynein light and light intermediate chains, but not with LIS1. LIS1 interacts with motor domains and the stalk.
